# Modulation of *TTR* Gene Expression in the Eye using Modified Duplex RNAs

**DOI:** 10.1101/2025.03.11.642595

**Authors:** Jiaxin Hu, Xin Gong, Jayanta Kundu, Dhrubajyoti Datta, Martin Egli, Muthiah Manoharan, V. Vinod Mootha, David R. Corey

## Abstract

Small interfering RNAs (siRNAs) are a proven therapeutic approach for controlling gene expression in the liver. Expanding the clinical potential of RNA interference (RNAi) requires developing strategies to enhance delivery to extra-hepatic tissues. In this study we examine inhibiting *transthyretin* (*TRR*) gene expression by short interfering RNAs (siRNAs) in the eye. Anti-*TTR* siRNAs have been developed as successful drugs to treat TTR amyloidosis. When administered systemically, anti-*TTR* siRNAs alleviate symptoms by blocking *TTR* expression in the liver. However, TTR amyloidosis also affects the eye, suggesting a need for reducing ocular *TTR* gene expression. Here, we demonstrate that C5 and 2’-O-linked lipid-modified siRNAs formulated in saline can inhibit *TTR* expression in the eye when administered locally by intravitreal (IVT) injection. Modeling suggests that length and accessibility of the lipid chains contributes to *in vivo* silencing. GalNAc modified anti-dsRNAs also inhibit *TTR* expression, albeit less potently. These data support lipid modified siRNAs as an approach to treating the ocular consequences of TTR amyloidosis. Inhibition of *TTR* expression throughout the eye demonstrates that lipid-siRNA conjugates have the potential to be a versatile platform for ocular drug discovery.

**Graphical Abstract:** 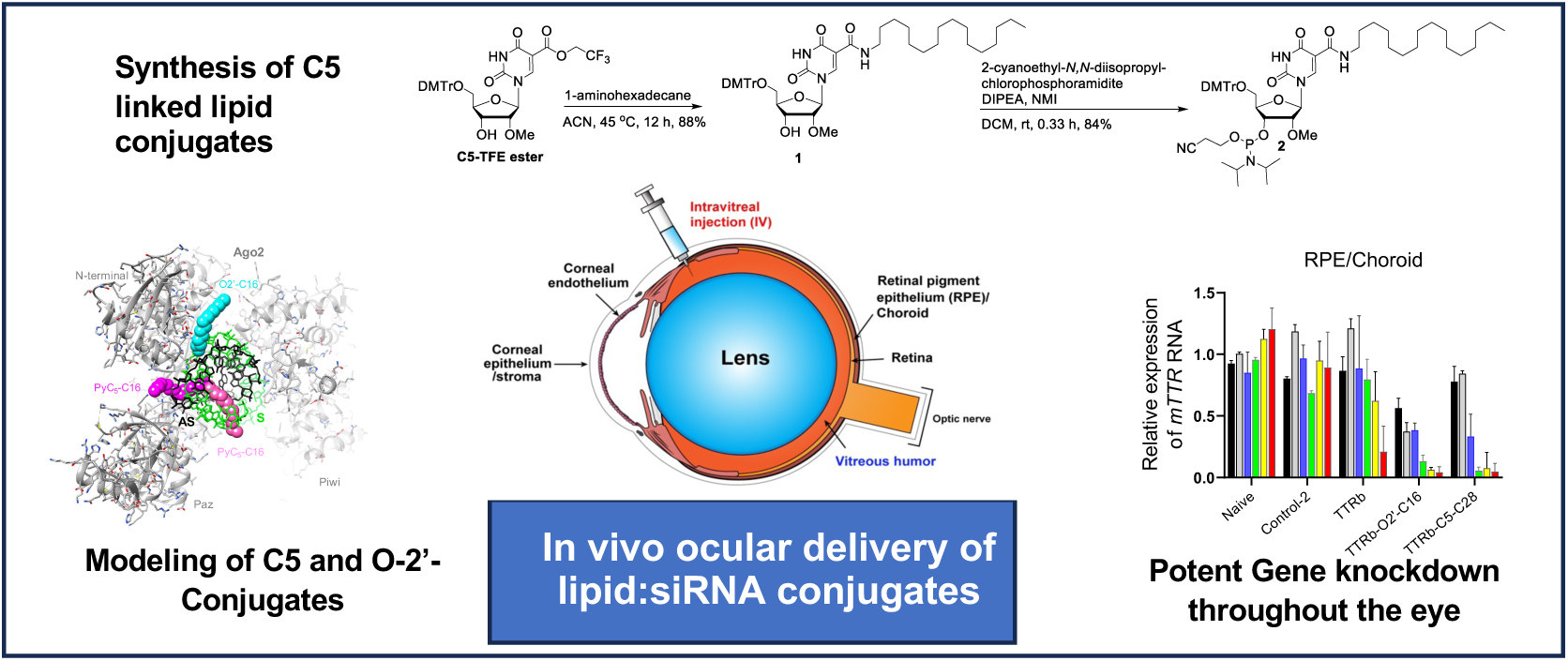

## Introduction

Small interfering RNAs (siRNAs) that act through RNA interference (RNAi) are a powerful approach for drug discovery and development (1–3). There are currently six approved siRNA drugs, all silencing mRNA targets in the liver. These positive clinical results and associated pre-clinical results demonstrate the power of siRNAs to be versatile regulators of gene expression in hepatic tissue. The open question is whether potent, selective, and clinically effective RNAi-mediated gene silencing can be extended to extra-hepatic tissues.

The success of siRNAs in the liver was enabled by the development of siRNA-GalNAc conjugates (4). GalNAc conjugates target the asialoglycoprotein receptor (ASGPR) and dramatically increase uptake of siRNAs in hepatocytes. Increased uptake leads to high potencies and ultimately contributes to greater success in the clinic. GalNAc conjugates are successful because of the large number of ASGPRs per cell. Unfortunately, the identification of receptors on other tissues that share the favorable properties of ASGPR has been slow, leading to interest in other modification strategies for optimizing siRNA delivery (1–3).

Strategies for extra-hepatic delivery have included cholesterol modification to improve skeletal delivery (5,6), antibody conjugates that improve delivery to muscle (7), and divalent siRNAs that improve delivery to the central nervous system (CNS) (8). Recently, lipid modification has been described as a general approach to delivery to extra-hepatic tissues, including CNS, lung, and eye (9).

The eye is a potentially advantageous organ for the delivery of nucleic acid drugs. The first approved nucleic acid drug, the antisense oligonucleotide (ASO) Vitravene (fomivirsen) in the 1990’s was administered by (IVT) injection (10). The second approved nucleic acid drug, the aptamer macugen, was also administered by IVT injection (11). Injection into the eye reduces the amount of drug needed and lowers the potential for systemic toxicities. IVT injection is one of the most common procedures in medicine and silencing gene expression in the eye might have therapeutic applications to many ocular diseases.

Our goal here was to explore the use of chemically modified lipid siRNA conjugates throughout varied parts of the eye. We chose *transthyretin* (*TTR*) as our target gene. *TTR* is the target of two approved siRNA drugs, one delivered using nanoparticles (12) and the other as a GalNAc conjugate (13). Systemic administration is sufficient to target *TTR* in the liver and alleviate many of the devastating symptoms of TTR amyloidosis (ATTR). ATTR can also lead to loss of vision by causing vitreous opacities, glaucoma, keratoconjunctivitis sicca (dry eye), and neurotrophic corneal disease (14–17). Systemic administration of the approved siRNA drugs does not treat ocular systems, presumably because of the barriers between the bloodstream and intraocular tissue (18). *TTR* expression, therefore, serves as a model gene for evaluating the action of siRNAs in the eye.

While ninety percent of TTR protein is synthesized by the liver, TTR is also produced by the choroid plexus in the brain and retinal pigment epithelial (RPE) cells in the eye (19). Progression of ocular involvement of ATTR is observed in patients even after liver transplantation (a surgical treatment for ATTR), despite the inability of plasma TTR to cross the blood-retina barrier (16,20). This finding suggests that ocular symptoms of ATTR are due to the local production of mutant TTR in the RPE.

Intraocular amyloid deposits have been detected in tissue structures across the eye including the corneal endothelium, trabecular meshwork, iris epithelium, ciliary pigment epithelium, lens capsule, vitreous, RPE, and retinal nerve fibers (15,17,21). ATTR can result in deposits at the inner pupillary margin resulting in its scalloped appearance and an altered pupillary light reflex (18). The reported prevalence of vitreous opacities in ATTR ranges from 5.4 to 35% (17). The collagen in the vitreous acts as a scaffold for the accumulation of amyloid that can result in a visual complaint of floaters and progressive loss of visual acuity.

The current surgical treatment to restore vision is vitrectomy but the opacities may recur. Approximately 18% of patients can also develop a chronic open angle glaucoma due to amyloid deposition in the trabecular meshwork and/or increased episcleral venous pressure (17). The glaucoma may be refractory to topical glaucoma medications and require surgery. These deficiencies in current treatments suggest a need for reduction of ocular TTR gene expression.

Here we show that lipid modified siRNAs (**Figure 1, Supplemental Table S1**) are potent inhibitors of *TTR* expression throughout diverse tissue in the eye, suggesting that lipidated siRNAs may be a broadly useful starting point for ocular drug discovery beyond their potential use limiting *TTR* expression. Molecular modeling suggests that the length of the lipid chain and its point of attachment are critical variables governing activity. Specific to ATTR, GalNAc modified siRNAs are less efficacious but also inhibit *TTR* expression, suggesting that drugs that are already approved for systemic administration might be candidates for IVT administration to patients at risk for ocular sequelae from ATTR. Potent inhibition by multiple lipidated anti-*TTR* siRNAs suggests that lipid conjugates can also provide strategy for treating the ocular findings of ATTR.

**Figure 1.**
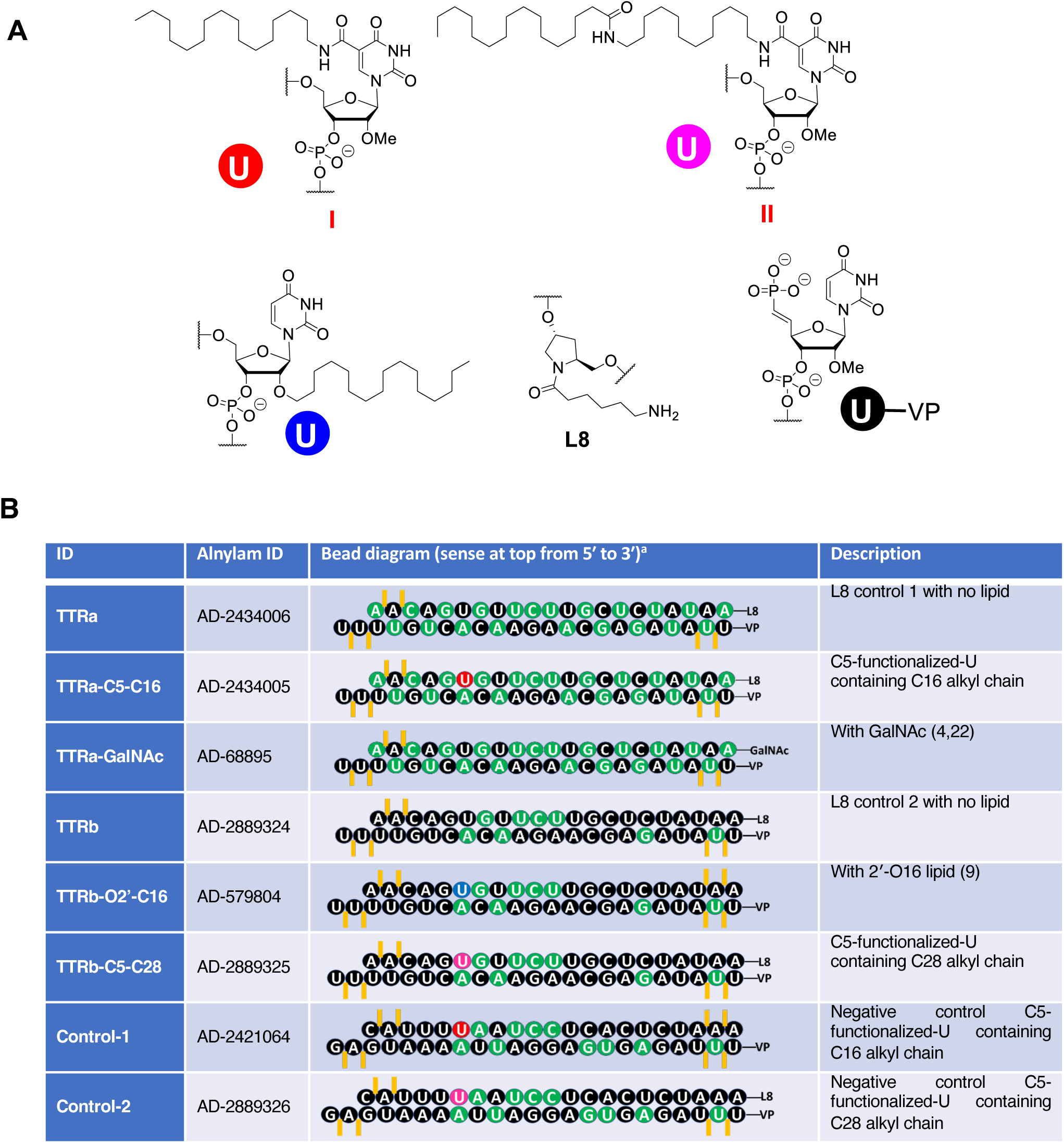
(A) Structural representation of C5-lipid, L8 linker, and vinyl phosphonate (VP) modifications used in lipid modified siRNAs. (B) Duplexes used in these studies. 2′-fluoro (2′-F) and 2′-*O*-methyl (2′-OMe) nucleotides are indicated in green and black, respectively. Phosphorothioate (PS) linkages are indicated by orange lines. VP refers to vinyl phosphonate. Uridines linked to lipid at the C5 position are red, uridines linked to lipid at the 2’-O position are blue.

## Materials and Methods

### General Synthetic Protocols

TLC was performed on Merck silica gel 60 plates coated with F254. Compounds were visualized under UV light (254 nm) or after spraying with the p-anisaldehyde staining solution followed by heating. Flash column chromatography was performed using a Teledyne ISCO Combi Flash system with pre-packed RediSep Teledyne ISCO silica gel cartridges. All moisture-sensitive reactions were carried out under anhydrous conditions using dry glassware, anhydrous solvents, and argon atmosphere. All commercially available reagents and solvents were purchased from Sigma-Aldrich unless otherwise stated and were used as received. ^1^H NMR spectra were recorded at 600 MHz (**Supplemental Figure S4**). ^13^C NMR spectra were recorded at 151 MHz. ^31^P NMR spectra were recorded at 243 MHz. Chemical shifts are given in ppm referenced to the solvent residual peak (CDCl_3_ – ^1^H: δ at 7.26 ppm and ^13^C δ at 77.2 ppm). Coupling constants are given in Hertz. Signal splitting patterns are described as singlet (s), doublet (d), triplet (t) or multiplet (m).

### 1-((2R,3R,4R,5R)-5-((bis(4-methoxyphenyl)(phenyl)methoxy)methyl)-4-hydroxy-3-methoxytetrahydrofuran-2-yl)-N-hexadecyl-2,4-dioxo-1,2,3,4-tetrahydropyrimidine-5-carboxamide *1*

To a clear solution of 2,2,2-trifluoroethyl 1-((2R,3R,4R,5R)-5-((bis(4-methoxyphenyl)(phenyl)methoxy)methyl)-4-hydroxy-3-methoxytetrahydrofuran-2-yl)-2,4-dioxo-1,2,3,4-tetrahydropyrimidine-5-carboxylate (C5-TFE ester) (2) (1.0 g, 1.46 mmol) in acetonitrile (ACN) (10 mL) was added 1-aminohexadecane (1.06 g, 4.38 mmol) and the reaction mixture was allowed to stir at 45 °C for 12 h. TLC showed that starting material was almost consumed (90 %). The volatile matter of the reaction mixture was concentrated under high vacuum pump to get the crude compound which was purified by flash column chromatography eluting with 30-50 % EtOAc in hexane to afford **1** (1.05 mg, 88% yield) as yellowish white solid. ^1^H NMR (600 MHz, CDCl_3_) δ 8.56 – 8.44 (m, 2H), 7.47 – 7.43 (m, 2H), 7.38 – 7.33 (m, 4H), 7.29 – 7.25 (m, 2H), 7.20 – 7.16 (m, 1H), 6.87 – 6.78 (m, 4H), 5.88 (s, 1H), 4.11 (d, *J* = 6.2 Hz, 1H), 4.05 – 4.01 (m, 1H), 3.87 (dd, *J* = 5.8, 3.1 Hz, 1H), 3.78 (s, 6H), 3.53 (s, 3H), 3.45 (t, *J* = 4.7 Hz, 2H), 3.37 – 3.32 (m, 2H), 1.55 (d, *J* = 7.4 Hz, 2H), 1.25 (s, 26H), 0.88 (t, *J* = 7.0 Hz, 3H) ppm. ^13^C NMR (151 MHz, CDCl_3_) δ 160.9, 160.9, 158.5, 149.2, 146.2, 144.7, 135.8, 135.8, 130.1, 130.1, 128.1, 127.9, 126.8, 113.2, 113.2, 107.0, 89.2, 86.7, 83.6, 82.8, 69.8, 63.3, 58.9, 55.2, 39.5, 31.9, 29.7, 29.7, 29.7, 29.7, 29.6, 29.5, 29.4, 29.4, 27.1, 22.7, 14.1 ppm.

### (2R,3R,4R,5R)-2-((bis(4-methoxyphenyl)(phenyl)methoxy)methyl)-5-(5-(hexadecylcarbamoyl)-2,4-dioxo-3,4-dihydropyrimidin-1(2H)-yl)-4-methoxytetrahydrofuran-3-yl (2-cyanoethyl) diisopropylphosphoramidite *2*

To a clear solution of **1** (1.02 g, 1.25 mmol) in anhydrous dichloromethane (DCM) (15 mL) were added *N*-methylimidazole (NMI) (204.95 mg, 2.50 mmol, 198.98 μL) and *N,N*-diisopropylethylamine (DIPEA) (806.54 mg, 6.24 mmol, 1.09 mL). After stirring for 5 minutes, 2-cyanoethyl-*N,N*-diisopropylchlorophosphoramidite (590.8 mg, 2.50 mmol, 556.8 μL) was added and the reaction mixture was allowed to stir for 0.33 h. TLC showed complete consumption of the starting material. The reaction mixture was quenched with aq NaHCO_3_, diluted with anhydrous dichloromethane and washed with water and brine. The organic layer was dried over Na_2_SO_4_, filtered and concentrated to get the crude product which was purified by silica gel column chromatography eluting with 40-60 % EtOAc-hexane to afford **2** (1.06 g, 84 % yield) as a white solid. ^1^H NMR (600 MHz, CDCl_3_) δ 8.59 – 8.42 (m, 2H), 7.46 (ddt, *J* = 13.8, 6.5, 1.4 Hz, 2H), 7.40 – 7.32 (m, 4H), 7.30 – 7.25 (m, 2H), 7.19 (qd, *J* = 6.1, 1.3 Hz, 1H), 6.91 – 6.77 (m, 4H), 5.96 – 5.86 (m, 1H), 4.30 – 4.17 (m, 2H), 4.02 – 3.93 (m, 1H), 3.79 (d, *J* = 4.4 Hz, 6H), 3.68 – 3.50 (m, 3H), 3.44 (d, *J* = 10.8 Hz, 3H), 3.36 (ddd, *J* = 14.8, 5.5, 2.7 Hz, 4H), 2.67 – 2.29 (m, 2H), 1.56 (q, *J* = 6.6 Hz, 2H), 1.25 (s, 24H), 1.15 (dd, *J* = 6.8, 5.3 Hz, 9H), 1.03 (d, *J* = 6.8 Hz, 4H), 0.88 (t, *J* = 7.0 Hz, 3H) ppm. ^13^C NMR (151 MHz, CDCl_3_) δ 162.4, 160.9, 158.5, 149.2, 146.4, 146.3, 144.6, 135.8, 135.7, 135.6, 130.2, 130.2, 130.1, 128.2, 128.1, 127.9, 126.8, 117.8, 113.2, 113.2, 113.2, 113.2, 107.0, 106.9, 88.9, 88.8, 86.7, 86.7, 83.5, 83.5, 83.4, 83.3, 82.2, 82.1, 81.4, 81.4, 71.0, 70.9, 70.5, 70.4, 63.2, 62.8, 58.7, 58.7, 58.6, 58.3, 58.3, 57.9, 57.8, 55.2, 55.2, 43.4, 43.3, 43.2, 43.1, 39.5, 39.4, 31.9, 29.7, 29.7, 29.7, 29.6, 29.6, 29.5, 29.5, 29.4, 27.1, 24.6, 24.6, 24.6, 24.5, 24.5, 22.7, 20.4, 20.3, 20.2, 20.1, 14.1 ppm. ^31^P NMR (243 MHz, CDCl_3_) δ 150.95, 150.20 ppm.

### N-(12-aminododecyl)-1-((2R,3R,4R,5R)-5-((bis(4-methoxyphenyl)(phenyl)methoxy)methyl)-4-hydroxy-3-methoxytetrahydrofuran-2-yl)-2,4-dioxo-1,2,3,4-tetrahydropyrimidine-5-carboxamide *3*

To a clear solution of C5-TFE ester(21) (1 g, 1.46 mmol) in anhydrous ACN (15 mL) was added dodecanediamine (291.80 mg, 1.46 mmol) and the reaction mixture was heated at 50 °C for 12 h. TLC showed complete consumption of the starting material. The volatile matter of the reaction mixture was concentrated under high vacuum pump to get the crude product which was purified by silica gel column chromatography eluting with 0-30 % MeOH-DCM to afford **3** (922 mg, 80 % yield) as white foam. ^1^H NMR (600 MHz, CDCl_3_) δ 8.70 (t, *J* = 5.6 Hz, 1H), 8.47 (s, 1H), 7.48 – 7.42 (m, 2H), 7.37 – 7.32 (m, 4H), 7.28 – 7.23 (m, 3H), 7.21 – 7.16 (m, 1H), 6.86 – 6.80 (m, 4H), 5.89 (d, *J* = 3.1 Hz, 1H), 4.23 (s, 4H), 4.09 (dd, *J* = 7.0, 5.7 Hz, 1H), 4.02 (ddd, *J* = 7.0, 5.6, 3.8 Hz, 1H), 3.85 (dd, *J* = 5.8, 3.1 Hz, 1H), 3.77 (s, 6H), 3.51 (s, 3H), 3.47 – 3.41 (m, 2H), 3.38 – 3.32 (m, 2H), 2.74 (t, *J* = 7.2 Hz, 2H), 1.55 (t, *J* = 7.3 Hz, 2H), 1.49 (t, *J* = 7.1 Hz, 2H), 1.35 – 1.24 (m, 16H) ppm. ^13^C NMR (151 MHz, CDCl_3_) δ 164.0, 161.5, 158.6, 150.4, 145.9, 144.9, 136.0, 135.9, 130.3, 130.2, 128.3, 128.0, 126.9, 113.3, 113.3, 107.2, 89.1, 86.7, 83.6, 83.0, 70.0, 63.6, 58.9, 55.3, 41.5, 39.3, 32.5, 29.4, 29.3, 29.2, 29.2, 29.1, 29.1, 29.0, 26.8, 26.7 ppm.

### 1-((2R,3R,4R,5R)-5-((bis(4-methoxyphenyl)(phenyl)methoxy)methyl)-4-hydroxy-3-methoxytetrahydrofuran-2-yl)-2,4-dioxo-N-(12-palmitamidododecyl)-1,2,3,4-tetrahydropyrimidine-5-carboxamide *4*

To a clear solution of **3** (850 mg, 1.08 mmol) in anhydrous DCM (15mL) were added 2,5-dioxopyrrolidin-1-yl palmitate (458.18 mg, 1.30 mmol) and DIPEA (279.20 mg, 2.16 mmol, 376.27 μL) and the reaction mixture was allowed to stir at rt for 12 h. TLC showed complete consumption of the starting material. The volatile matters of the reaction mixture was concentrated under high vacuum pump to get the crude product which was purified by silica gel column chromatography eluting with 20-50 % EtOAc-hexane to afford **4** (1.03 g, 93 % yield) as white foam.^1^H NMR (600 MHz, CDCl_3_) δ 9.90 (d, *J* = 3.3 Hz, 1H), 8.41 – 8.31 (m, 2H), 7.25 (d, *J* = 7.8 Hz, 2H), 7.16 (dd, *J* = 8.5, 5.3 Hz, 3H), 7.07 (dd, *J* = 10.5, 5.6 Hz, 3H), 6.98 (dd, *J* = 8.1, 5.6 Hz, 1H), 6.68 – 6.59 (m, 4H), 5.70 (d, *J* = 3.1 Hz, 1H), 5.41 (t, *J* = 5.7 Hz, 1H), 3.93 (q, *J* = 6.8 Hz, 1H), 3.85 (dt, *J* = 10.0, 4.7 Hz, 1H), 3.68 (dd, *J* = 5.8, 3.1 Hz, 1H), 3.58 (d, *J* = 7.0 Hz, 6H), 3.37 – 3.30 (m, 3H), 3.26 (tt, *J* = 10.8, 6.5 Hz, 2H), 3.16 (dq, *J* = 12.7, 6.8 Hz, 2H), 3.03 (q, *J* = 6.8 Hz, 2H), 2.67 (dd, *J* = 7.7, 3.1 Hz, 1H), 1.95 (t, *J* = 7.7 Hz, 2H), 1.47 – 1.25 (m, 7H), 1.07 (d, *J* = 14.0 Hz, 42H), 0.68 (t, *J* = 6.9 Hz, 3H) ppm. ^13^C NMR (151 MHz, CDCl_3_) δ 173.3, 162.9, 161.2, 158.5, 149.5, 146.2, 144.8, 139.6, 135.9, 135.9, 130.2, 130.1, 129.2, 128.2, 127.9, 127.9, 127.8, 126.8, 113.3, 113.2, 113.2, 107.0, 89.2, 86.7, 83.6, 82.9, 69.8, 63.5, 58.9, 55.2, 39.6, 37.0, 32.0, 29.8, 29.7, 29.7, 29.6, 29.6, 29.5, 29.5, 29.4, 29.4, 29.3, 29.3, 27.1, 27.0, 25.9, 22.8, 14.2 ppm.

### (2R,3R,4R,5R)-2-((bis(4-methoxyphenyl)(phenyl)methoxy)methyl)-5-(2,4-dioxo-5-((12-palmitamidododecyl)carbamoyl)-3,4-dihydropyrimidin-1(2H)-yl)-4-methoxytetrahydrofuran-3-yl (2-cyanoethyl) diisopropylphosphoramidite *5*

To a clear solution of **4** (900 mg, 877.74 μmol) in anhydrous DCM (10 mL) were added DIPEA (567.20 mg, 4.39 mmol, 764.42 μL) and NMI (144.13 mg, 1.76 mmol, 139.80 μL). The reaction mixture was allowed to stir for 5 minutes and then 2-cyanoethyl-*N,N*-diisopropylchlorophosphoramidite (415.49 mg, 1.76 mmol) was added and the reaction mixture was left at rt for additional 0.33 h. TLC showed complete consumption of the starting material. The reaction mixture was quenched with saturated aq. NaHCO_3_ and diluted with DCM. The organic layer was dried over Na_2_SO_4_ and filtered. Solvent was removed in rotary evaporator to get the crude product which was purified by silica gel column chromatography eluting with 20-40 % EtOAc-hexane to afford **5** (973 mg, 90 % yield) as white foam.^1^H NMR (600 MHz, CDCl_3_) δ 9.61 (s, 1H), 8.58 – 8.48 (m, 2H), 7.49 – 7.41 (m, 2H), 7.40 – 7.32 (m, 4H), 7.30 – 7.24 (m, 2H), 7.21 – 7.15 (m, 1H), 6.86 – 6.80 (m, 4H), 5.97 – 5.89 (m, 1H), 5.62 – 5.46 (m, 1H), 4.31 – 4.16 (m, 2H), 4.02 – 3.92 (m, 1H), 3.86 (ddd, *J* = 7.5, 6.0, 2.9 Hz, 2H), 3.79 – 3.76 (m, 6H), 3.55 (tdd, *J* = 16.9, 7.9, 5.3 Hz, 2H), 3.44 (d, *J* = 9.2 Hz, 3H), 3.39 – 3.31 (m, 3H), 3.22 (q, *J* = 6.7 Hz, 2H), 2.61 (q, *J* = 6.4 Hz, 1H), 2.34 (td, *J* = 6.4, 4.6 Hz, 1H), 2.14 (t, *J* = 7.6 Hz, 2H), 1.65 – 1.52 (m, 4H), 1.47 (t, *J* = 7.1 Hz, 2H), 1.37 – 1.19 (m, 43H), 1.15 (t, *J* = 6.3 Hz, 8H), 1.01 (d, *J* = 6.7 Hz, 4H), 0.87 (t, *J* = 7.0 Hz, 3H) ppm. ^13^C NMR (151 MHz, CDCl_3_) δ 173.3, 162.8, 161.2, 158.6, 158.5, 149.6, 146.3, 144.8, 144.7, 135.9, 135.7, 135.7, 130.3, 130.2, 130.2, 128.3, 128.2, 127.9, 126.8, 117.9, 117.6, 113.3, 113.3, 113.3, 113.2, 107.1, 107.0, 88.9, 88.8, 86.7, 86.7, 83.5, 83.4, 83.3, 82.3, 81.6, 71.1, 71.0, 70.6, 70.5, 63.3, 62.9, 58.8, 58.8, 58.8, 58.7, 58.4, 58.3, 58.0, 57.9, 55.2, 43.5, 43.4, 43.3, 43.2, 39.6, 39.5, 37.0, 32.0, 29.8, 29.8, 29.7, 29.7, 29.6, 29.6, 29.6, 29.5, 29.4, 29.4, 29.3, 27.1, 27.0, 25.9, 24.7, 24.6, 24.6, 22.8, 20.4, 20.4, 20.2, 20.2, 14.2 ppm.^31^P NMR (243 MHz, CDCl_3_) δ 150.89, 150.20 ppm.

### Oligonucleotide synthesis and purification

Oligonucleotides were synthesized on K & A Labs H-8 automated oligonucleotide synthesizer at 10 μmol scale using suitable supports. A solution of 0.25 M 5-(*S*-ethylthio)-1*H-*tetrazole in ACN (ACN) was used as the activator. The solutions of commercially available phosphoramidites and synthesized phosphoramidities were used at 0.15 M in anhydrous ACN. The oxidizing reagent was 0.02 M I_2_ in THF/pyridine/H_2_O. *N,N*-Dimethyl-N′-(3-thioxo-3H-1,2,4-dithiazol-5-yl)methanimidamide (DDTT) in 0.1 M in pyridine was used as the sulfurizing reagent. The detritylation reagent was 3% dichloroacetic acid in DCM. Detritylation of modified building blocks was performed manually using 3 % TCA-DCM. Waiting times for coupling, capping, oxidation, and sulfurization step were 450 s, 25 s, 80 s, and 300 s, respectively. Lipid modified building blocks were coupled manually with two syringes and coupling time was 15 min. After completion of the automated synthesis, the oligonucleotide was manually released from support and deprotected using Ammonium Hydroxide/40% aqueous MethylAmine 1:1 v/v (AMA) at rt for 3 h.

After filtration through a 0.45-μm nylon filter, oligonucleotides were quantified. For ion exchange, a preparative HPLC column custom packed with TSKgel SuperQ-5PW (20) (Sigma) was used. Appropriate gradients of mobile phase (buffer A: 20 mM sodium phosphate, 15% ACN, pH 8.5; buffer B: 1 M NaBr, 20 mM sodium phosphate, 15% ACN, pH 8.5) were employed. Oligonucleotides were desalted using size-exclusion chromatography using three Hi-Prep columns connected in series and water as an eluent. Oligonucleotides containing lipids were purified in RP-HPLC using appropriate gradient of buffer A (3 % ACN in 50 mM aq triethylammonium acetate) and buffer B (80 % ACN in 50 mM aq triethylammonium acetate) at ambient temperature (**Supplementary Figure S4**). Pure fractions were concentrated in rotary evaporator at 36 °C and desalted by size-exclusion chromatography using three Hi-Prep columns connected in series and water as an eluent. For sodium exchange, lyophilized powders were dissolved in 0.1 M sodium acetate and desalted again following the same desalting process. Oligonucleotides were then quantified by measuring the absorbance at 260 nm.

Extinction coefficients were calculated using the following extinction coefficients for each residue: A, 13.86; T/U, 7.92; C, 6.57; and g, 10.53 M^-1^cm^-1^. The identities of modified oligonucleotides were verified by mass spectrometry (**Table S1**). Purities were evaluated by analytical reverse-phase HPLC. For reverse-phase HPLC, a C-18 column was used with a gradient of 2-45 % buffer B (buffer A: 95 mM hexafluoroisopropanol, 16.3 mM TEA, 0.05 mM EDTA; buffer B: MeOH) over 39 min. All the IEX-HPLC were performed in DNAPac PA200 BioLC (4×250 mm) analytical column. Buffer A is 15 % ACN, 20 mM sodium phosphate (pH 11) and buffer B is 15 % ACN, 20 mM sodium phosphate, 1 M NaBr (pH 11). A gradient of 31-60 % B over 12 min is used at 30 °C. See **Table 1** for sequences of C5-lipid modified oligonucleotides used for this study.

### Measuring thermal stability for C5 modifications by UV-T_m_

siRNA duplexes were annealed by heating at 94 °C for 2 min followed by slow cooling (23,24). For T_m_ measurement, all the duplexes were used at 1 µM concentration in 0.1XPBS. To minimize the possibility of any secondary structures cooling curve was recorded first by cooling the duplexes from 95 °C to 25 °C at a cooling rate of 1 °C/min. For heating curve, the duplexes were heated from 25 °C to 95 °C at a heating rate of 1 °C/min. For both cooling and heating curves data was recorded in 1 min interval. The final T_m_ is the average of 3 heating curves and 3 cooling curves which has been provided in table.

### HPLC-based hydrophobicity assay for C5 lipids containing duplexes and single strands

Retention time (RT) from non-denaturing ion-pairing reverse-phase high-performance liquid chromatography (nd IP-RP HPLC) was used as a measure of duplex hydrophobicity. Duplexes were injected into C8 column (Agilent Poroshell 120 EC-C8: 2.1×100 mm, 2.7 µm, 120 Å) and eluted with suitable gradient using mobile phase A (100 mM TEAA) and B (100 mM TEAA in 85% MeOH). For single strands hydrophobicity, C5-modification containing sense strands and controls were injected into C8 column (Waters XBridge® BEH C8 2.5μM 2.1X50mm) and eluted with suitable gradient using mobile phase A (95 mM hexafluoroisopropanol, 16.3 mM triethylamine, 50 µM EDTA) and B (MeOH).

### Intraocular injections in mice

All animal experiments were approved by the Institutional Animal Care and Use Committee of University of Texas Southwestern Medical Center. The C57BL6J mice were purchased from the University of Texas Southwestern Mouse Breeding Core and equal number of age-matched litter-mate male and female mice were used when possible. Mice were given unlimited access to water and food and were on a twelve-hour light/twelve-hour dark cycle.

Mice were anesthetized using intraperitoneal injections of ketamine/xylazine (120 mg/kg, 16 mg/kg, respectively) followed by dilation of pupils with one drop of phenylephrine 1 % (Alcon) and 1 drop of tropicamide 2.5 % (Alcon) 2 minutes afterwards. Upon full dilation, lubricant ointment eye gel (Henry Schein) was applied to the ocular surface to prevent desiccation of the corneas. The IVT injections were performed as previously described and guided using a Zeiss microscope(23).

The anti-*TTR* siRNAs were pre-formulated with 1X Phosphate-buffered saline (PBS) to reach the desired concentration. A 30-gauge beveled needle was introduced either one mm posterior to the supero-lateral limbus at a 45-degree angle toward the vitreous cavity for IVT injection. The needle was inserted 1.5 mm deep into the eye and then carefully removed. The egressed chamber fluid was removed using a cotton tip applicator. A Hamilton micro-syringe fitted to a 33-gauge beveled needle was placed in the previously fashioned needle tract and the solution was slowly injected in the eye. The needle was held in place for an additional minute after complete injection to allow cavity fluid equilibration with the injected solution. Post-injection AK-polysporin-bacitracin antibiotic ointment (Perrigo) was applied to the needle wound followed by a drop of the lubricant gel. The mice were kept on a heating pad while recovering from the anesthesia.

Subcutaneous Injections were administered into the loose skin over the mouse neck.

### Ocular tissue harvest

Given the small amount of tissue available after dissection of the mouse eye, analysis of *TTR* mRNA levels by qPCR is challenges and requires strict adherence to protocols. Following euthanization of the mouse, whole globes were enucleated at the optic nerve, rinsed with balanced salt solution (BSS), and then submerged in a small droplet of BSS in a 100 mm x 15 mm plastic petri dish. The globes were then punctured at the corneal limbus with a 30-gauge needle and circumferentially bisected along the limbus to separate the anterior and posterior segments. The neurosensory retinal tissue from the posterior chamber was carefully separated from the underlying retinal pigment epithelium (RPE), choroid, and scleral tissue. These tissue layers were rinsed with BSS before storage. Whole corneal tissue from the anterior chamber was preserved after removal of lens and pigmented iris tissue. Lens capsule was peeled free and rinsed with BSS before storage.

Proceeding whole mouse corneal acquisition, the endothelial surface of the corneal tissue was incubated in trypan blue stain (Stephens Instruments) for two minutes before rinsed and re-submerged in BSS, allowing for enhanced visualization of the corneal endothelium apparent by the appearance of a thin translucent blue layer within the concave surface of the cupped cornea. Next, a 30-guage needle was used to score the perimeter of the corneal endothelium. Then, forceps were used to grasp the stained endothelium edge followed by gentle separation of the thin blue endothelial layer from the underlying corneal stroma and epithelium. These tissue layers were then rinsed with BSS before storage.

All collected ocular tissues were inspected under high power microscopy for any signs of congenital, pathological or procedure-related defects such as cataracts, opacities, fibrosis or obvious signs of edema, which may indicate congenital malformations, procedure-related trauma or endophthalmitis, or abnormal wound healing. Any tissue with apparent defects were discarded and not used for quantitative analysis. Harvested corneal epithelium/stroma, corneal endothelium, lens capsule epithelium, neurosensory-retina, and RPE/choroid/scleral tissues together with liver tissue for siRNA-GalNAc were used to assess efficacy of TTR expression knockdown.

### RNA extraction and qPCR analysis of TTR expression

Total RNA from corneal endothelium and lens capsule epithelium was extracted using RNeasy Micro Kit (Qiagen), RNA from corneal epithelium/stroma, retina, and RPE/choroid were extracted by RNeasy Mini kit (Qiagen). Tissues were lysed using a homogenizer for 30 seconds to 1 minute with interval, other steps were followed according to the manufacturer’s protocol. cDNAs were made by reverse transcription with high-capacity reverse transcription kit (Applied Biosystems) per the manufacturer’s protocol using equal amount of RNAs from tissue samples. *q*PCR experiments were performed on a CFX96 Touch real-time PCR system (Bio-Rad) using iTaq SYBR Green Supermix (Bio-Rad). PCR reactions were done in triplicates at 55°C 2 min, 95°C 3 min and 95°C 30 s, 60°C 30 s for 40 cycles in an optical 96-well plate. Data was normalized relative to levels of reference gene *RPL19* mRNA. Mouse *TTR* primers: F, 5’- CGTACTGGAAGACACTTGGCATT; R, 5’-GAGTCGTTGGCTGTGAAAACC. Mouse RPL19 primers: F 5’- GTATGCTCAGGCTACAGAAGAG-3’; R 5’- GAGTTGGCATTGGCGATTT-3’. The final data were analyzed using GraphPad Prism 8 software.

## RESULTS

### Lipid conjugate design and synthesis

Duplex RNAs were synthesized with a mixture of 2′-fluoro (2′-F) and 2′-*O*-methyl (2′-*O*-me) bases to improve *in vivo* stability and activity (**Figure 1, Supplemental Table S1**). Phosphorothioate linkages (PS) or an L8 linker were included at termini to improve resistance to degradation by nucleases.

The length and attachment point of the lipid chain can affect binding to soluble proteins and membranes, affecting the biodistribution of nucleic acids. C16 lipid chains were attached via the C5 position of a nucleobase (TTRa-C5-C16) or the O2′ position of the ribose (TTRb-O2’-C16). TTRb-C5-C28 had a C28 lipid chain conjugated to the C5 position of the nucleobase.

TTRa-GalNAc has the same sequence as TTRa-C5-C16, but the GalNAc is attached at the 3′ end of sense strand with a hydroxyprolinol linker. TTRa-GalNAc is similar in design to the approved drug Vutrisiran (13). TTRa and TTRb lack the conjugated lipid but have similar chemical modifications to TTRa-C5-C16 and TTRb-O2’-C16/TTRb-C5-C28 respectively. Control-1 and Control-2 have similar arrangements of chemical modifications as TTRb-O2’-C16/TTRb-C5-C28 but lack complementarity to TTR mRNA.

For compound TTRa-C5-C16, we first synthesized hexadecyl lipid chain containing 2′-*O*-methyluridine phosphoramidite (**Figure 2A, Supplemental Figure S1**) where the lipid chain was present at C5-position of the nucleobase. Activated trifluoro ester (C5-TFE-ester) (22) was heated with 1-aminohexadecane to get compound **1** via amide bond formation. Compound **1** was then employed for phosphoramidite synthesis using standard protocol to produce lipophilic phosphoramidite **2** in good yield which was incorporated in sense strands of siRNA using standard oligonucleotides synthesis protocols.

**Figure 2.**
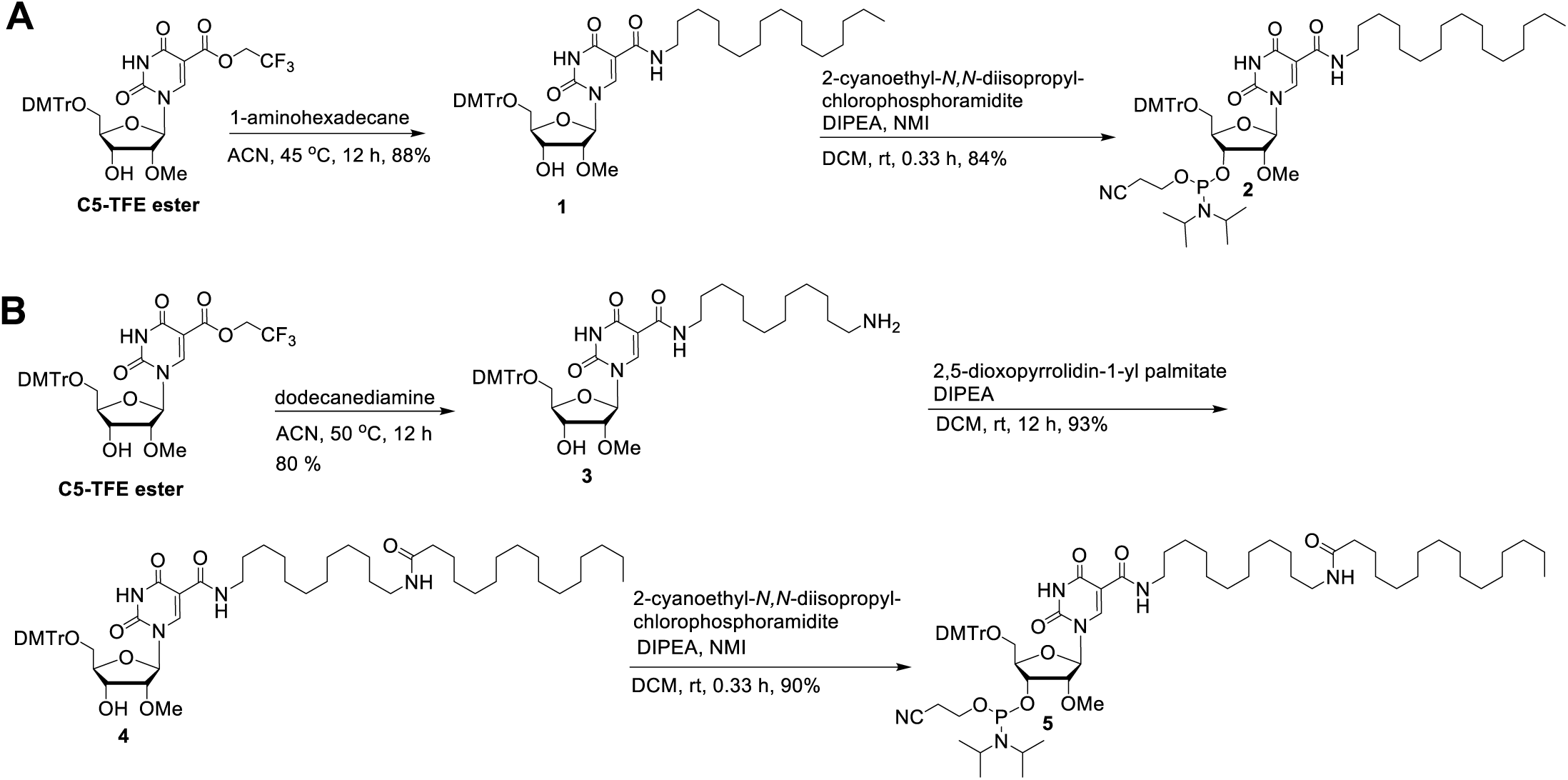
(A) Synthesis of C5 C16-lipid chain containing 2′-*O*-methyluridine phosphoramidite. (B) Synthesis of C5 C28-lipid chain containing 2′-*O*-methyluridine phosphoramidite synthesis.

With compound TTRb-C5-C28, we explored the role of longer lipid chain by synthesizing C28-lipid containing phosphoramidite (**Figure 2B**) (22–24). First, activated trifluoro ester (C5-TFE-ester) was heated with 1,12-diaminododecane to obtain compound **3** via amide bond formation. Compound **3** was then reacted with commercially available N-hydroxysuccinimide ester of hexadecenoic acid for a second amide bond formation to have compound **4** which was finally employed for phosphoramidite synthesis using standard protocol to have lipophilic phosphoramidite **5** in good yield.

### Effect of C5-lipid attachment in the context of duplex relative to single strands

Hydrophobicity is one of the critical determinants of biodistribution. To evaluate the hydrophobicity contribution from C5-lipids attached to single strands relative to duplexes we used reverse phase HPLC (**Supplemental Figure S2**). As expected, both TTRb-C16 and TTRb-C28 were more hydrophobic than TTRb regardless of whether the modified strand was single-stranded or in a duplex. C28 conjugation had a bigger impact than attachment of C16, as would be expected from the longer C28 lipid chain.

The impact for C28 relative to C16, however, was much greater in the single stranded context. For single strands, C28-lipid at C5 position of uridine showed almost 5 minutes delay in elution (13.5 min) as compared to standard 2′-O-C16 lipid (8.5 min). In duplexes, by contrast, C28-lipidated duplex TTRb-C5-C28 showed only 1.3 min elution delay (11.3 min) with respect to 2′-O-C16 lipid (10.0 min) duplex TTRb-O2’-C16. One explanation for the muted impact of C28 on hydrophobicity of lipidated duplexes is that the lipid is less able to bind into the major groove when attached at C5 than when attached at O2’, making interactions with the reverse phase matrix more likely.

Thermal melting studies revealed that duplex RNA TTRb-C5-C28 is thermally destabilized by 8.0 degrees as compared to control (no lipid) duplex TTRb whereas TTRb-O2’-C16 is only 1.5 degrees destabilized relative to TTRb (**Supplementary Figure S3, Supplementary Table S2**). This is consistent with the lipid component of TTRb-C5-C28 having fewer stabilizing interactions with the RNA duplex. Both steric and electrostatic factors may contribute to the greater loss in stability caused by the longer lipid chain. It is more challenging to accommodate the C28 inside the major groove, a site of limited space and strong negative electrostatic potential. Any conformation of this chain that does not eject a C12 portion out of the cavity will interfere with hydration and pairing. The shorter C16 chain has fewer opportunities to interfere with pairing and nucleobase stacking interactions. Even when the C28 chain manages to partially protrude from the groove, it might move around the backbones of the duplex, interfere with phosphate and sugar hydration, and obstruct metal ion binding.

### *In vitro* evaluation of dsRNA conjugates

Attachment of lipid or GalNAc moieties to a siRNA might affect the intrinsic silencing potency of the duplex, independent of any positive effects on biodistribution. To test whether this was occurring, we transfected unmodified control duplexes TTRa and TTRb into mouse BNL CL2 liver cell line and compared their activities to modified duplexes TTRa-C5-C16, TTRb-O2’-C16, TTRb-C5-C28, and TTRa-GalNAc.

Spontaneous uptake of lipid-modified siRNAs is not efficient in cultures cells and was facilitated using the cationic lipid RNAiMAX.

Attachment of C16 at C5 to create TTRa-C5-C16 had little effect on potency when evaluated relative to the unconjugated duplex TTRa in cultured cells (**Figure 3a**). IC_50_ values were approximately 1-3 nM regardless of whether the lipid was attached. Unmodified duplex TTRb was much less potent than TTRa, indicating that the chemical modification pattern used for the TTRb series of compounds reduces potency. This deficit, however, was rescued by attachment of lipid as both TTRb-O2’-C16 and TTRb-C5-C28 were like the potency of TTRa-C5-C16 (**Figure 3b**). This surprising result may indicate that the chemical modification pattern used in the TTRb series reduces *in vitro* potency which is then rescued by attachment of lipid. These data suggest that lipid modified RNA duplexes have good potencies inside cells in culture and supported further testing in mice.

**Figure. 3.**
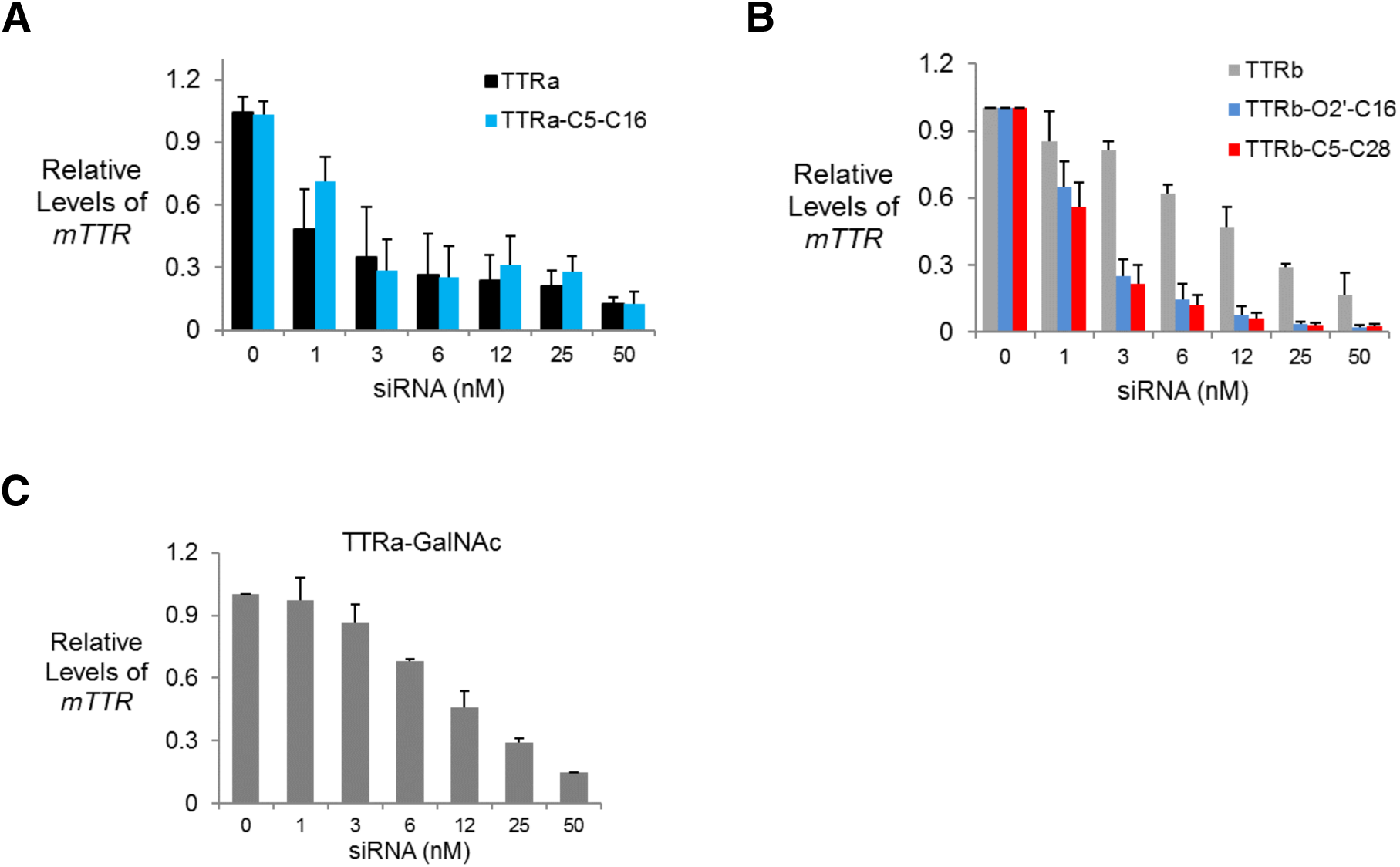
*TTR* gene expression after transfection of lipid modified dsRNAs into BNL CL2 murine cells using RNAiMAX as a transfection reagent. (A) TTRa and TTRa-C5-C16. (B) TTRb, TTRb-O-2’-C16, and TTRb-C5-C26. (C) TTRa-GalNAc. Error bars are mean with SEM. N=4 independent replicates.

We also evaluated the GalNAc to 3’ terminus of the passenger strand because to evaluate whether conjugates that are like the approved drug Vutrisiran. We observed that the attachment of GalNAc to TTRa reduced potency in cell culture with an IC_50_ value of approximately 12 nM (**Figure 3c**). GalNAc conjugation, despite immense value for the success of in vivo drug development, may be at a slight disadvantage relative to lipid conjugates for fundamental intracellular silencing.

### Gene silencing throughout the eye

Our goal was to analyze tissues from both the anterior and posterior segments of the mouse eye including the retinal pigment epithelium (RPE) (**Figure 4A**). The eye is a complex organ composed of tissues with diverse functions that offer different drug targets and opportunities of oligonucleotide drug development. It is important, therefore, to analyze the different parts of the eye separately. In the mouse eye, this approach is challenged by the small amounts of critical tissues and the associated need to micro-dissect samples.

**Figure. 4.**
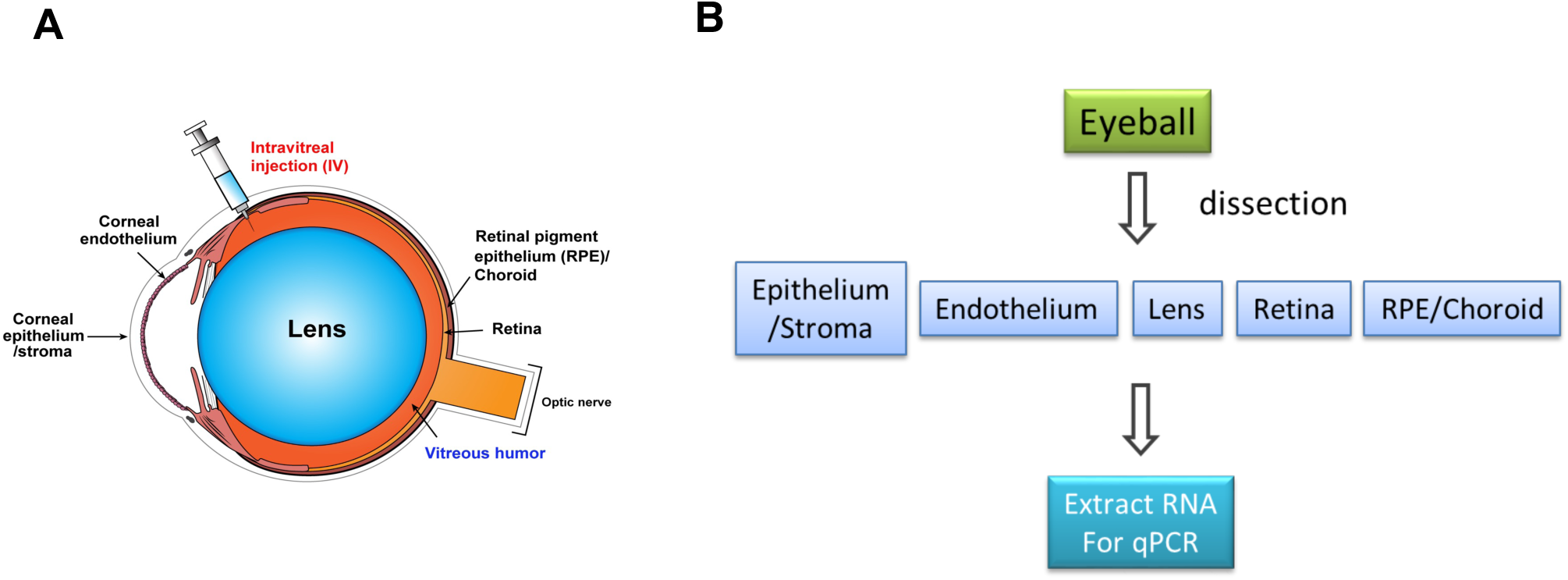
Strategy for ocular injections and analysis. (A) Schematic for the murine eye. (B) Scheme for dissection and tissue analysis. Tissue was harvested seven days after intravitreal (IVT) injection.

While we examined TTR expression throughout eye, the RPE is the primary tissue involved with ocular ATTR. The RPE is a continuous monolayer of post-mitotic cells located between the neurosensory retina and the choroid, the vascular connective tissue layer that nourishes the outer retina. The lateral membranes of the RPE cells are joined by a continuous belt of tight junctions that form the outer blood-retina barrier (25). The RPE is metabolically active in its maintenance role of photoreceptor layer by regulating the flow of nutrients and waste products to and from the retina and renewal of the spent outer segments of the photoreceptors. Ocular production of mutant TTR by the RPE is thought to result in ATTR eye disease findings (16,17,25).

siRNAs were formulated in PBS and administered by intravitreal (IVT) injection to test their effects on *TTR* expression. We had previously observed that delivery of anti-MALAT1 ASOs by IVT was more effective than delivery by intracameral injection (IC) into the anterior chamber (26,27). Tissue was harvested from the corneal epithelium/stroma, corneal endothelium, the lens, the neurosensory retina, and the RPE/choroid layers (**Figure 4B**). We processed these tissues and evaluated expression using quantitative PCR (qPCR).

Prior to testing conjugates in mice, we evaluated the relative expression of *TTR* mRNA in different eye tissues (**Figure 5**). *TTR* was expressed most highly in the RPE, consistent with the previous observation that cultured RPE cells synthesize and secrete the TTR protein(19,28). Expression of *TTR* mRNA was 40-fold higher in RPE/choroid than retina and 40- to 10,000-fold higher than in lens, corneal endothelium, or corneal epithelium/stroma.

**Figure 5.**
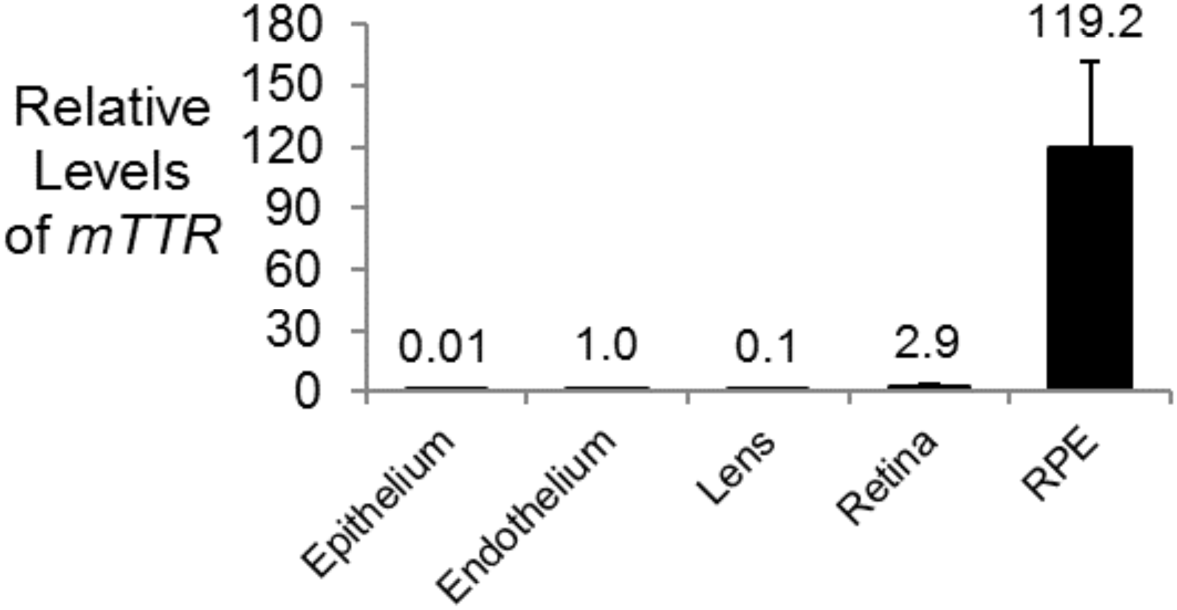
Relative expression of *TTR* mRNA in different murine ocular tissues measured by qPCR. Data was normalized relative to levels of murine *RPL19*. Error bars are mean with SEM. N=4 independent replicates.

### Testing of C5 linked C16 conjugate in mice administered by intravitreal injection

We began by analyzing the effect of siRNA conjugate TTRa-C5-C16 on *TTR* expression in mouse RPE/choroid, lens, corneal endothelium, corneal epithelium/stroma, and retina (**Figure 6**). Two mice were injected with 50 µg compound per eye for each experimental group. (**Figure 6A-E**). For all tissues, greater than 80% inhibition was achieved relative to eyes injected with PBS or Control-1 duplex RNA. This initial testing demonstrated that siRNA conjugates had the potential to inhibit gene expression in all parts of the mouse eye. We then examined inhibition as a function of concentration of conjugates TTRa and TTRa-C5-C16 in the RPE. Inhibition of *TTR* expression was not efficient at when less than 9 µg was injected, suggesting a need for more potent designs for ocular use of lipid-conjugated siRNAs. (**Figure 6F,G**).

**Figure. 6.**
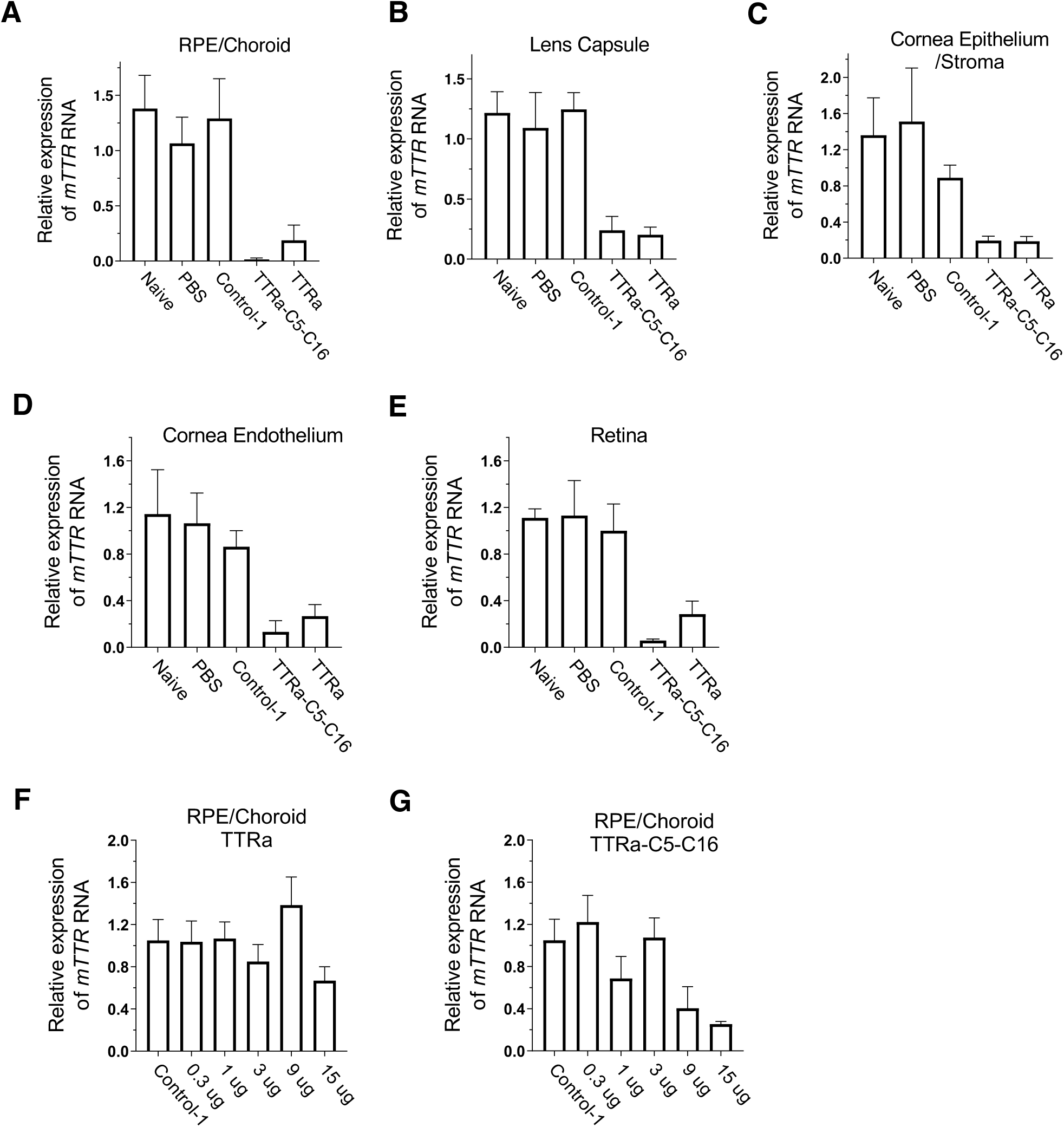
Inhibition of *TTR* gene expression by dsRNAs administered (IVT) at 50 µg per eye. (A) RPE/choroid. (B) Lens capsule. (C) Corneal epithelium/stroma (D) Corneal endothelium. (E) Retina. (F,G) Dose response for TTRa and TTRa-C5-C16. Tissue was harvested seven days after injection. Data was normalized relative to levels of murine *RPL19*. Error bars are mean with SEM. N=4 independent replicates.

### Testing of C5 linked C28 and 2’-O C16 conjugate in mice

We reasoned that altering the point of lipid attachment or the length of the lipid chain might enhance potency. To test this hypothesis, we evaluated RNA duplexes TTRb, 2’O linked conjugate TTRb-O2’-C16, and C5 linked conjugate TTRb-C5-C28 (**Figure 7**). As further described below, modeling suggests that both TTRb-O2’-C16 and TTRb-C5-C28 have improved potential to form protein interactions that might affect biodistribution and delivery to cells. These duplexes also have a different pattern of modifications than TTRa-C5-C16 that has been observed to offer improved potencies against other targets. This new generation of compounds was chosen for detailed dose response analysis over a range of concentrations.

**Figure. 7.**
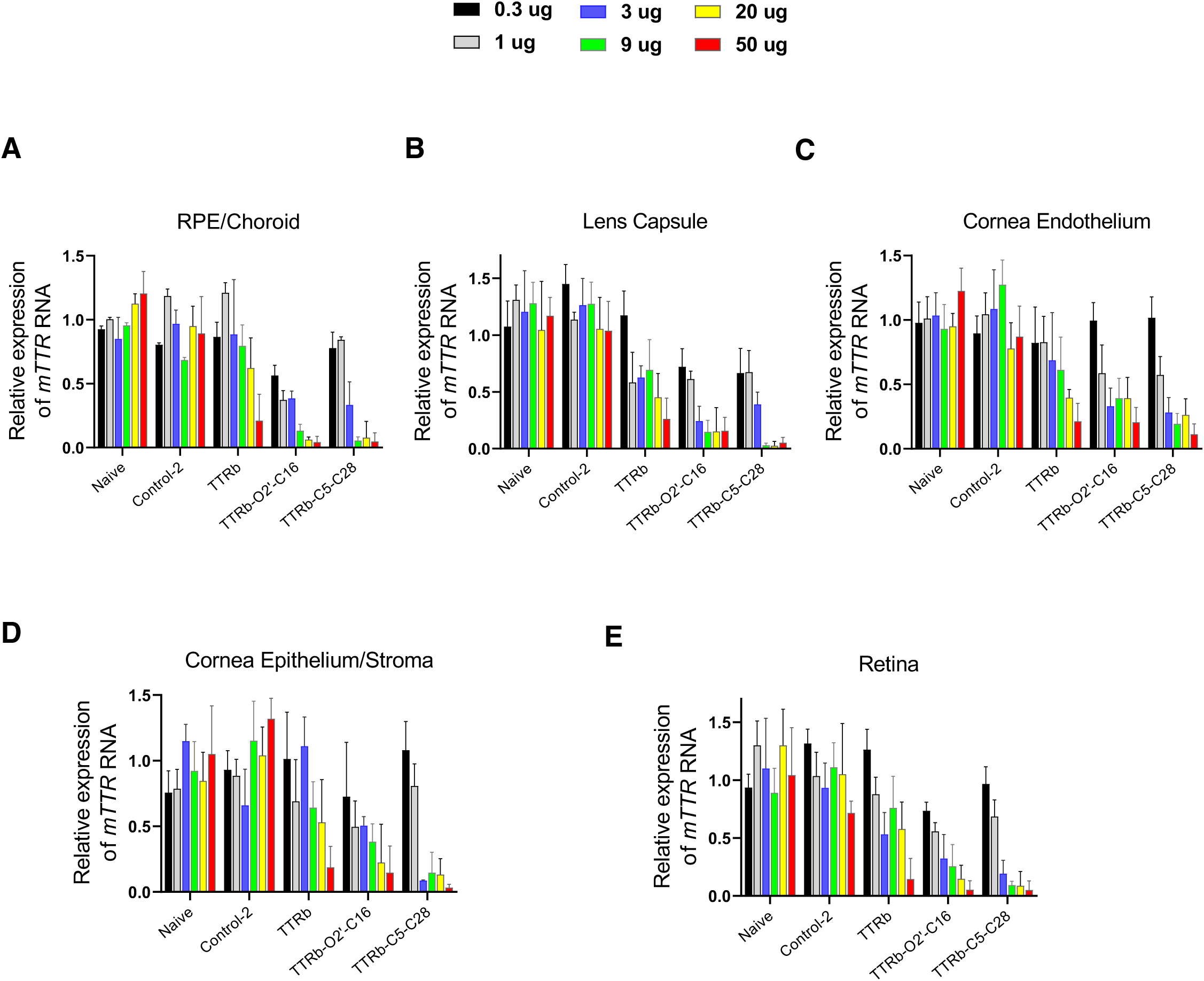
Inhibition of *TTR* gene expression in the RPE by dsRNAs TTRb-C5-C16, TTRb-O2’-C16, and TTRb-C5-C28 at varying concentrations. Tissue was harvested seven days after IVT injection. Data was normalized relative to levels of murine *RPL19*. Error bars are mean with SEM. N=4 independent replicates.

We observed that TTRb, the siRNA that lacked lipid modification, inhibited *TTR* expression but only at the highest dose (50 µg) in RPE/choroid (**Figure 7A**). By contrast, conjugates TTRb-O2’-C16 and TTRb-C5-C28 were more potent inhibitors, with greater than 90% inhibition when 9 µg were injected and ∼50% inhibition after injection of as little as 0.3-1 µg. Duplex RNA Control-2 had no significant effect on RPE/choroid expression, even when administered at the highest dose. TTRb-C5-C28 achieves potent gene silencing even though thermal melting studies reveal that it has a significantly lower T_m_ than TTR or TTRb-O2’-C16 (**Supplemental Table 2, Supplemental Figure 2**).

While the RPE is an important source of production of the mutant TTR protein responsible for ATTR ocular pathology, we also examined the effect of siRNAs on *TTR* expression in other parts of the eye, including lens, corneal endothelium, corneal epithelium/stroma, and retina (**Figure 7B-E**). Measuring the effects of dsRNA-modulated *TTR* gene expression in these tissues of the eye is more challenging because of the low levels of *TTR* RNA relative to RPE/choroid. While there was greater uncertainty in the data, we observed clear trends showing the potential of lipid-modified siRNAs to broadly inhibit gene expression in tissues of both the anterior and posterior segment of the eye.

### Inhibition of TTR expression by GalNAc conjugates

A GalNAc conjugate, TTRa-GalNAc (**Figure 1**), is like the conjugate siRNA already approved to treat ATTR. Our goal was to test whether it could also inhibit TTR expression in the eye. TTRa-GalNAc was administered either systemically by subcutaneous (SubQ) injection or directly into the eye by IVT injection.

When administered SubQ, TTRa-GalNAc inhibited *TTR* gene expression in the liver at ∼90% (**Figure 8A**), consistent with the known hepatic efficacy of GalNAc conjugates. *TTR* expression after SubQ administration was not inhibited in the RPE, consistent with the bloodstream eye barrier (24) (**Figure 8B**). When TTRa-GalNAc was injected into the eye by IVT at 50 µg, inhibition of *TTR* mRNA expression was observed (**Figure 8B**). IVT injection of TTRa-GalNAc had no effect on *TTR* expression in the liver (**Figure 8A**), consistent with the presence of an ocular/blood barrier and the small amount of compound injected into the eye.

**Figure. 8.**
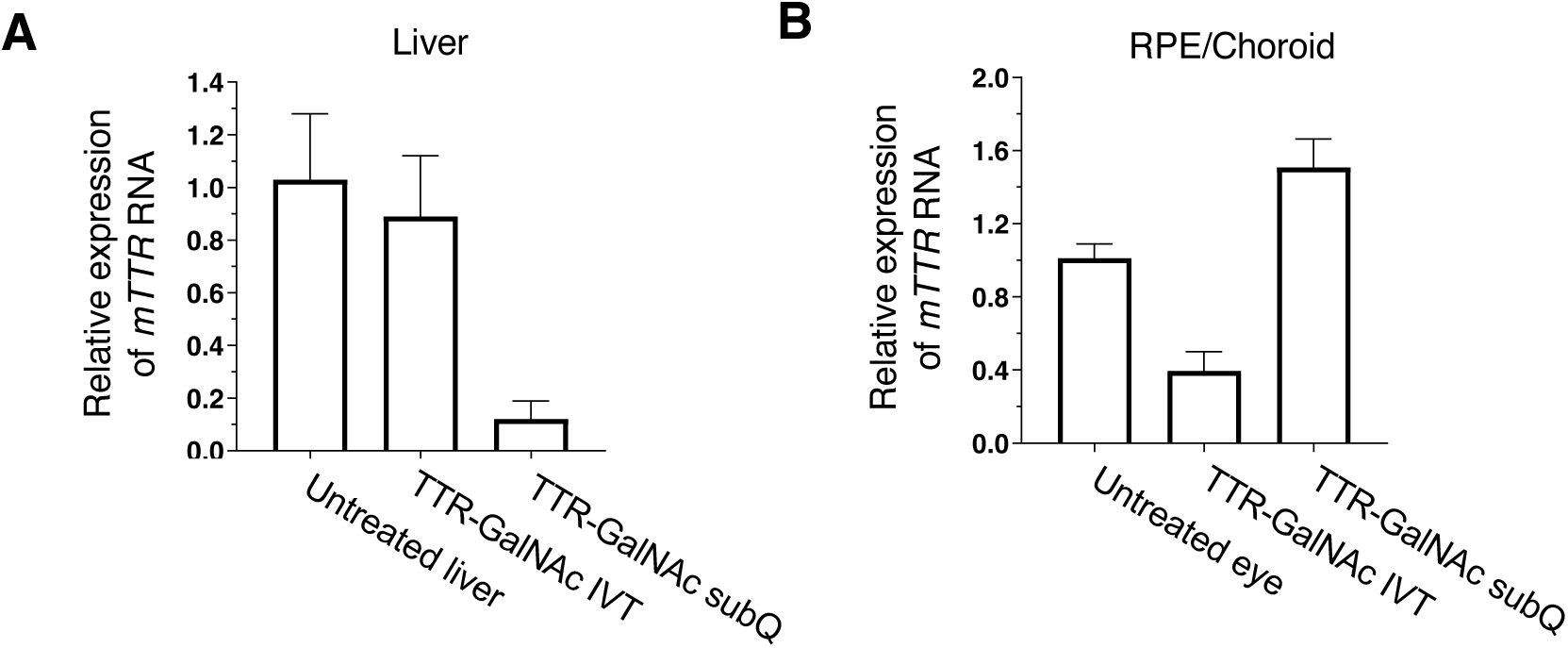
Inhibition of *TTR* gene expression in the RPE by dsRNA TTR-GalNAc in mouse RPE and liver. (A) Expression levels of *TTR* in mouse liver. (B) Expression levels in RPE/Choroid. Tissue was harvested seven days after injection of 3 µg (IVT) or 50 µg (SubQ). Data was normalized relative to levels of murine *RPL19*. Error bars are mean with SEM. N=4 independent replicates.

### Modeling of the binding of human serum albumin to siRNAs with different lipid modifications at the C-5 position

TTRa-C5-C16 and TTRb-C5-C28 share a common attachment for the lipid chain to the C5 position of uridine 6 of the sense strand (**Figure 1**). While TTRb-C5-C28 was a potent *in vivo* silencing agent for *TTR* gene expression (**Figure 7**), TTRa-C5-C16 was less potent (**Figure 6G**). These data suggest that the longer lipid affects the efficacy of lipidated siRNAs. To gain insight into how the longer lipid or point of attachment might contribute to the different activities of siRNAs tested in this report, TTRa-C5-C16, TTRb-O2’-C16, and TTRb-C5’-C28 (**Figure 1**), we turned to human serum albumin (HSA) as a model system for protein-lipid interactions.

HSA interacts with synthetic oligonucleotides and plays an important role governing the systemic biodistribution of therapeutic nucleic acids (29), HSA is also the dominant soluble protein in vitreous fluid (30), suggesting that it has the potential to form interactions that impact the biodistribution and activity of therapeutic oligonucleotides in the eye.

To evaluate the accessibility of lipid chains for interactions with HSA, we examined the crystal structure of HSA in complex with myristic acid (C14) (**Supplementary Figure S5**) determined at a resolution of 1.9 Å that shows eight bound fatty acid molecules (PDB ID 8RCP) (31). We chose one of the myristic acid molecules with the carboxylate located near the protein surface. Using the UCSF Chimera graphic software package (32), we converted the carboxylate C to methylene and extended the chain to C16. We first modeled C16 attached to the 2’ oxygen. The TTR siRNA model was generated in 3DNA (33) and then combined with the modified serum albumin-myristic acid complex to build the TTRa-O2’-C16 conjugate bound to albumin.

The lipid chain, now extended from C14 to C16, slightly protrudes from the albumin surface (**Figure 9A**). Using UCSF Chimera, the siRNA duplex together with torsion angles around the C13-C14 and C14-C15 bonds of the lipid were manipulated such that O2′ of sense strand U6 and the terminal methylene of the C16 chain were positioned for bond formation between the lipid and O2’ located in the shallow grove. The final model of TTRb-O2’-C16 does not exhibit unfavorable contacts between albumin and siRNA, all torsion angles along the C16 chain are in the ap or ac ranges, and the binding mode of the myristic acid portion of the C16 chain is that observed in the crystal structure of the complex (**Figure 9A**). This modeling suggests that the attachment of C16 to O2’ permits substantial interactions between lipid and HSA

**Figure 9.**
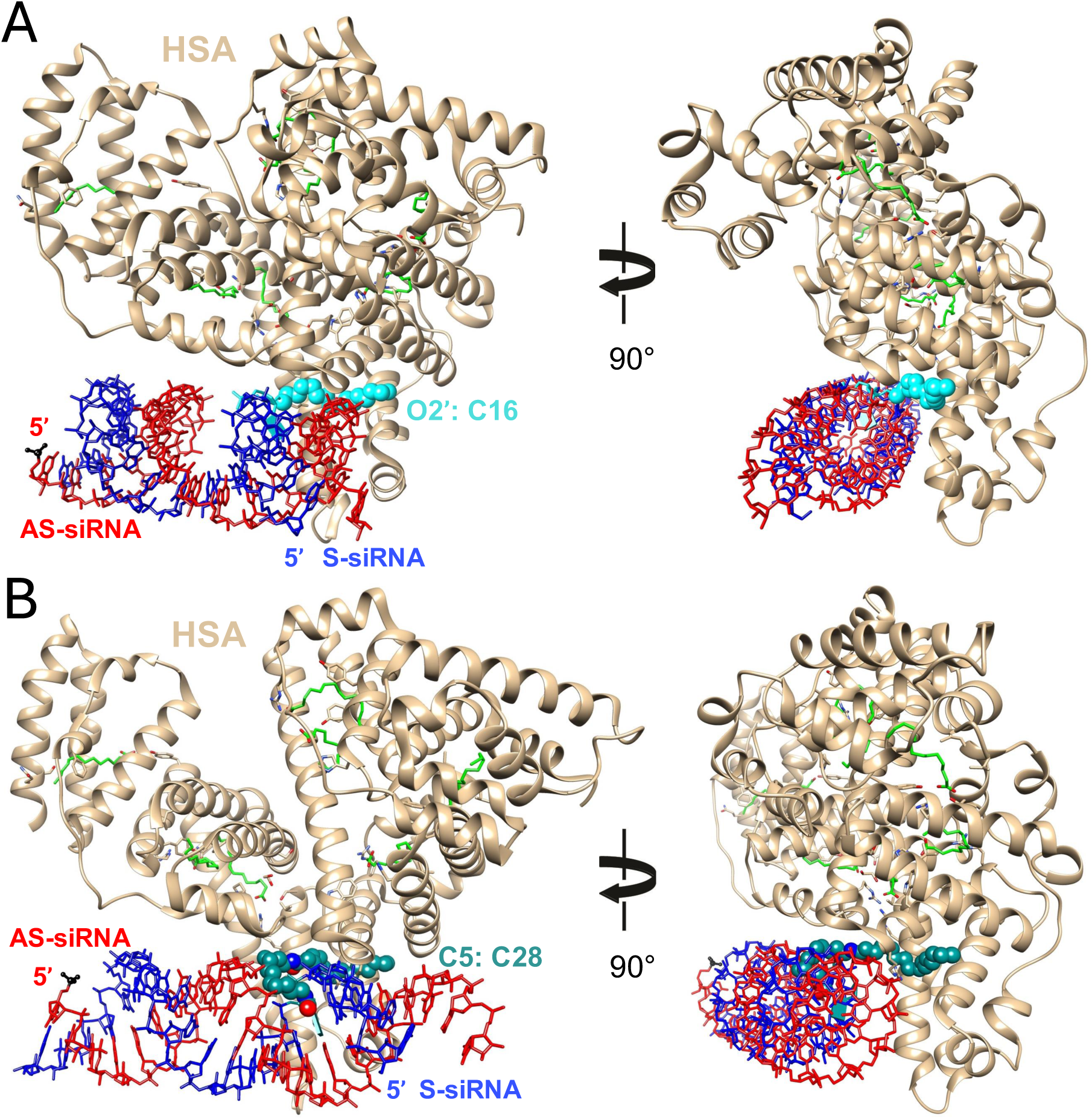
Computational models of the (A) TTRb-O2’-C16 and (B) TTRb-C5-C28 siRNA duplexes bound to human serum albumin based on the crystal structure of the albumin-myristic acid complex (PDB ID 8RCP). TTRa-C5-C16 can be imagined as a subset of the TTRb-C5-C28 model shown in (B). The views are across the major and minor grooves of the RNA duplex (left) and then rotated around the vertical by ca. 90 degrees and more along the helical axis (right). Albumin is depicted in cartoon mode and colored in tan and carbon atoms of fatty acid molecules are green. Bonds of siRNA antisense and sense strands are colored in red and blue, respectively, the 5′-terminal phosphate group of the former is highlighted in black and ball-and-stick mode (bottom left corner of figure panels on the left), and colors of C16 and C28 chain carbon atoms match those in Figure 1A.

We next evaluated C16 and C28 carbon chains attached at the C5 position. While we do not show the C16 chain separately, it is can be visualized as a subset of the C28 chain. We observed that the C16 chain is too short to link the lipid bound to albumin to the C5 position of U6 that is located at the floor of the narrow and deep major groove (**Figure 9B**). This helps rationalize the lack of activity seen with TTRa-C5-C16 – it lacks sufficient potential to escape the deep and narrow groove of the A-form siRNA duplex and form interactions with proteins like HSA.

The model we built of the TTRb-C5-C28 siRNA duplex bound to albumin is free of unfavorable contacts between RNA. In contrast to the C16 lipid, TTRb-C5-C28 has a lipid chain that is long enough to form an interface with HSA (**Figure 9B**). This model supports the hypothesis that the longer lipid chain allows greater binding to protein and may enhance potency *in vivo*. We note that this improved potency is achieved despite the intrinsic silencing of the non-lipid modified parent duplex of TTRb-C5-C28 being poorer than the intrinsic potency of TTRa-C5-C16 (**Figure 3**).

### Analysis of the binding of Argonaute 2 (Ago2) to siRNAs modified with C16 chains at O2’ or C5 positions

Next, we turned our attention to the interaction between siRNA-lipid conjugates and Argonaute2 (Ago2) endonuclease. Ago2 is the “slicer” enzyme that both promotes binding of siRNA guide strands to mRNA targets (32–34). Ago 2 is a critical member of the RNA-induced silencing complex (RISC) and cleaves RNA targets to amplify potency. Because of its central role in RNAi, interactions between Ago2 and the lipid chains may affect the potency of lipidated duplex RNA conjugates.

We modeled how attachment of C16 lipid at either the O2’ and C5 positions would affect association with Ago2. To build models of an siRNA duplex with O2’-C16 or C5-C16 conjugates bound to Ago2, we used the crystal structure of the enzyme in complex with miR-122 bound to a target RNA determined at a resolution of 3.4 Å(PDB ID 6MDZ)(31). We used UCSF Chimera to attach C16 chains to the O2′ and C5 positions and maintain C16 torsion angles in the ap and ac ranges. The C16 chain attached to O2′ in the shallow minor groove is almost completely exposed on the surface of the RNA duplex (**Figure 10**, cyan chain), allowing it to efficiently interact with lipid binding proteins and membranes.

**Figure 10.**
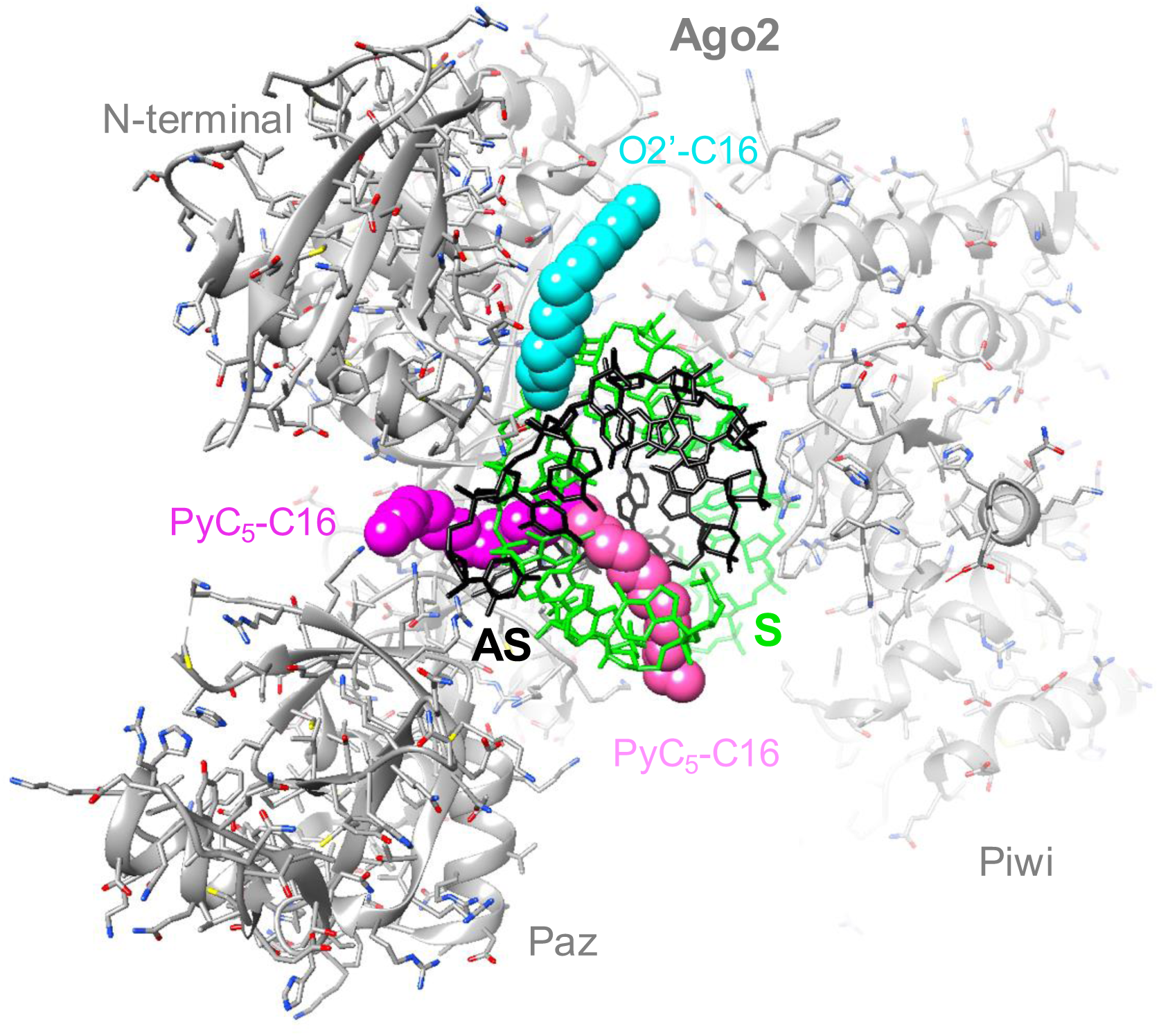
Computational models of human Ago2 in complex with an siRNA antisense (AS, black bonds)-sense (S, green bonds) strand duplex with C16 lipid conjugates attached to the O2′ or C5 positions of sense strand residue U6 based on the crystal structure of Ago2 bound to miR-122 opposite a target RNA (PDB ID 6MDZ). Ago2 is depicted in cartoon mode and colored in gray with individual domains labeled (the MID domain is invisible in the background). Carbon atoms of the C16 chain attached to O2′ are colored in cyan, The C16 chain attached to C5 was modeled in two alternative conformations, one emerging from the center of the major groove (carbon atoms colored in pink) and the other wrapping around the backbone of the antisense siRNA to escape the groove (carbon atoms colored in magenta).

Like our model of the TTRb-C5-C16 lipid siRNA conjugate bound to HSA (**Figure 9**), the C16 chain attached to C5 barely emerges from the deep major groove (**Figure 10**, pink and magenta chains adopt alternative conformations). Only four methylene moieties lie outside the groove, thereby illustrating that the C28 conjugate is needed to escape the major groove and present a sufficiently long lipid chain for stable interaction with molecules other than Ago2. The models of C16 chains (**Figure 9)** do not result in any short contacts to Ago2. If contacts beyond Ago2 contribute to the potency of lipidated C5 conjugates, C28 may have advantages versus C16.

## Discussion

### Expanding options for siRNA conjugates

Devising strategies for extra-hepatic delivery is the defining challenge for the development of nucleic acid therapeutics. The introduction of new conjugate types, lipids in this case, variations in the length of the chain, and the sites for attaching them help expand the repertoire of chemical tools for delivery to organs beyond liver. Potential beneficial effects include improvements in pharmacokinetics and pharmacodynamics, as well as better control of off-target effects. The exploration of new chemistries paves the road to gains in potency and safety, sometimes in ways that are not now evident.

### Ocular Efficacy of lipid conjugated siRNAs

Our primary goal was to test whether GalNAc and lipid-modified siRNAs could inhibit *TTR* gene expression in varied tissues within the mouse eye. A secondary goal was to explore how changing the length (C16 or C28) of the lipid chain or its point of attachment. (C5 or 2-O’) to the nucleotide would affect gene silencing. Our data from both C5 and 2’O-linked conjugates demonstrate inhibition in a broader range of ocular tissues, encouraging future exploration of gene silencing by lipid-modified siRNAs in tissues beyond the RPE. C16 and C28 conjugates tested after IVT injection inhibited gene expression in both the anterior and posterior segments of the eye, including the RPE/choroid, retina, lens, corneal endothelium, and corneal epithelium/stroma. Inhibition in all tissues was achieved even though *TTR* expression varies by 10,000-fold in these tissues.

### Designing base-modified (C5-uridine) lipid conjugates

Previous work had examined attachment of a C16 lipid chain a the 2’-O position of uridine 6 (9). The 2’-hydroxyl group allows for particularly facile attachment of substituents and linkers but could become problematic in a regiospecific fashion if the particular residue or ribose were engaged in protein-RNA interactions. The chemical stability of the major groove nucleobase site of attachment could be beneficial compared to the 2’-OH. Moreover, for oligomers like morpholino or PNA that typically lack a 2’-OH or similar functionality, C5 becomes more important.

We hypothesized that accessibility of the lipid chain to interactions with proteins would affect biodistribution and activity. To explore this concept, we initially synthesized C16 lipid conjugates at C5 position of uridine from activated trifluoro ester (21) (**Figures 1,2**). Initial *in vivo* study revealed that C16 lipid conjugate at C5 position (TTRa-C5-C16) yielded only modest improvement in potency relative to the analogous non-lipidated control (TTRa) (**Figure 6**). Reverse phase HPLC based hydrophobicity assay revealed that the C5-C16 conjugation contributes less hydrophobicity to the siRNA duplex than it does to the corresponding single strand, suggesting that the duplex blocks interactions with the C5-conjugated lipid chain.

After these observations we explored the activities of conjugates containing a longer lipid at C5, TTRb-C5-C28, or a lipid conjugated at the 2’-O position (TTRb-O2’-C16). Modeling studies suggested C16 lipid attached at the C5 position form interactions with the major groove of the siRNA. These interactions would obstruct interactions between the lipid chains and proteins like HSA or Ago2 (**Figures 9 and 10**) and might contribute to reduced potency. Modeling also suggested that a longer C5-linked lipid chain (TTRb-C5-C28) or a 2’-O linked lipid (TTRb-O2’-C16) would be more accessible. For C5 conjugates, the modeling suggests that the lipid should be at least 24 carbons.

Our experimental assays support these suggestions. TTRb-C5-C28 achieves a similar potency to the 2’-O linked conjugate TTRb-O2’-C16. While we acknowledge that more work will be necessary to substantiate our hypotheses connecting conjugated design, structure, and function. Our data, however, showing robust silencing by TTRb-O2’-C16 and TTRb-C5-C28 (**Figure 7**), suggests that strategic choice of lipid structure and placement can improve gene silencing.

### Ocular efficacy of GalNAc conjugated siRNAs

dsRNA-GalNAc conjugates were designed to exploit recognition of the ASGPR to efficiently deliver dsRNAs to the liver (40). These conjugates have been remarkably successful, resulting in several drugs that have a strongly favorable impact on the treatment of patients. TTR is a relatively rare disease, with 5000-7000 cases diagnosed within the United States each year. Only a fraction of these patients report intraocular symptoms (14–17), so a simple path to the clinic that does not require extensive testing of a new compound might be the most viable.

Our data suggest that inhibition of *TTR* expression can be achieved with GalNAc siRNA conjugates. Potency, however, is low relative to lipid conjugates and the lower potency would need to be addressed prior to consideration of GalNAc siRNA conjugates. Nevertheless, our results suggest that the application of existing siRNA drugs to target *TTR* expression is plausible and suggest thoughtful consideration related to optimizing delivery, potency, and safety.

### Comparing Ocular Efficacy of ASOs and siRNA

ASOs and dsRNAs are competing approaches to gene silencing. Comparing the two approaches is not straightforward because a fair comparison would require independent development projects aimed at identifying the optimal siRNA and the optimal ASO respectively for a given target. Previously, we examined gapmer ASOs targeting *MALAT-1*, a nuclear noncoding RNA that is widely used as a surrogate for ASO target engagement (23,26). Comparing the data our ASO and siRNA conjugate studies is not straightforward because the target genes are different. In addition, their mRNA transcripts are in different cellular compartments (*TTR* in cytoplasm, *MALAT-1* in the nucleus). IC_50_ values are approximations due to the relatively large error bars from measurements in tissues where *TTR* expression is low. Nevertheless, our past and present results are suggestive. We also note that Garanto and coworkers have demonstrated that a variety of different ASO chemistries can be active in the retina and can penetrate different layers of the retina (41).

The anti-MALAT-1 ASO inhibited MALAT-1 with IC_50_ values of approximately 0.5 to 1 µg/injection for each eye tissue. The most potent anti-*TTR* conjugates used in our current study, TTRb-O2’-C16 and TTRb-C5-C28 have similar approximate IC_50_ values. Within the acknowledged limitations of the comparisons, ocular administration of siRNAs and ASOs appears to have similar potential for gene silencing. Both are viable platforms for development of nucleic acid therapeutics in the eye. Moving forward, inhibition by siRNAs or ASOs suggests that these modalities are the foundation for development paths towards the treatment of other ocular diseases, such as Fuchs’ endothelial corneal dystrophy (33).

## Conclusions

Ocular silencing of *TTR* expression demonstrates the potential of lipid-modified siRNAs to inhibit gene expression throughout diverse eye tissues. Inhibition can be achieved in both the anterior and posterior segments of the eye. While subsequent studies will be necessary to fully characterize and optimize both dsRNA chemistry and injection routes of delivery, the inhibition of *TTR* that we observe suggests that the approach will be a valuable option for therapeutic development programs that aim to use gene knock down strategies to alleviate diseases of the eye. Reducing the 2′-F content from TTRa to TTRb in siRNA and changing major groove-minor groove impact biological potency due to the change in metabolic stability which was reflected in the divergent activities of the TTRa-C5-C16 and TTRb-O2’-C16 and TTRb-C5-C28 siRNA-lipid conjugates. It is likely that more potent conjugates await discovery and that the eye will become an increasingly important target for nucleic acid therapeutics.

## Acknowledgements

This study was supported by grants R01EY022161 (VVM), and R35GM118103 (DRC) from the National Institutes of Health, Bethesda, MD, a Challenge Grant from Research to Prevent Blindness (RPB), a Core Grant for Vision Research (P30EY030413), and the Welch Foundation (I-2184 to DRC). VVM is the Paul T. Stoffel / Centex Professor in Clinical Care. DRC is the Rusty Kelley Professor of Biomedical Science

## COMPETING INTEREST STATEMENT

M. Manoharan, J. Kundu and D. Datta are employees of Alnylam Pharmaceuticals. V.V.M., D.R.C., and J.H. hold a patent related to the use of dsRNAs to treat Fuch’s endothelial corneal dystrophy.

