## Supplemental for "Modulation of *TTR* Gene Expression in the Eye using Modified Duplex RNAs"

**Table of Contents**


Oligonucleotide synthesis scheme **S-2**
Oligonucleotide sequences and characterization **S-2**
$T_m$ data and representative heating curves **S-3**
HPLC based hydrophobicity assay **S-4**
Representative $T_m$ curves. **S-5**
$^{1}H$, $^{13}C$, and $^{31}P$ NMR spectra for the new compounds **S-6**
HPLC profiles of oligonucleotides **S-18**
Crystal structure of human serum albumin in complex with myristic acid **S-24**


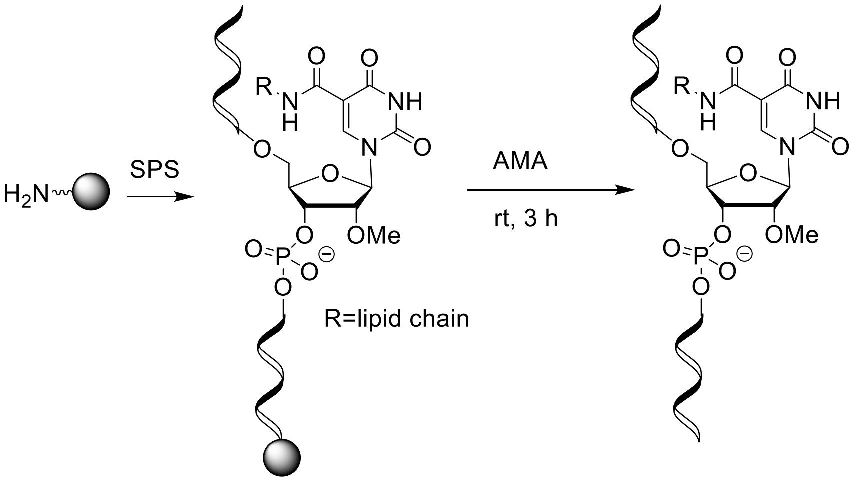


**Figure S1:** General synthesis scheme for oligonucleotides. SPS indicates solid-phase synthesis. AMA is (1:1) v/v 28-30 % aq. ammonia and 40 % aq. methylamine.

**Table S1:** Sequences and characterization of oligonucleotides used for this study.

| **Entry** | **Sequence of strands (5′-3′)** [a] | **Target** | **Mass(M-H)⁻** | |
|---|---|---|---|---|
| | | | **Calcd.** | **Obsd.** |
| ON1 | c●a●uuu**I***A*a*UCC*ucacucua●a●a | SOD1 | 7101.03 | 7100.43 |
| ON2 | c●a●uuu**II***A*a*UCC*ucacucuaaaL8 | SOD1 | 7558.50 | 7557.86 |
| ON3 | *A*●a●*C*a*G*u*G*u*UCU*u*G*c*U*c*U*a*U*a*A*L8 | TTR | 7093.48 | 7091.88 |
| ON4 | *A*●a●*C*a*G***I***G*u*UCU*u*G*c*U*c*U*a*U*a*A*L8 | TTR | 7360.94 | 7360.45 |
| ON5 | a●a●cagu*G*u*UCU*ugcucuauaaL8 | ocTTR | 7189.77 | 7188.92 |
| ON6 | a●a●cag**II***G*u*UCU*ugcucuauaaL8 | ocTTR | 7654.54 | 7653.82 |

[a]Chemical modifications are indicated as follows: ●, PS linkage; lower case, 2′-OMe; italicized upper case, 2′-F; L8, hydroxyprolinol-C6 linker.

**Table S2:** Melting temperatures of duplexes

| Entry | Bead diagram (sense at top from 5′ to 3′)[a] | Description | $T_m$[b](°C) | $\Delta T_m$[c](°C) |
|---|---|---|---|---|
| 1 | 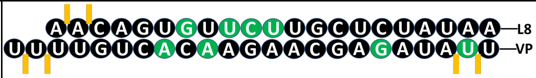 | L8 control 2 with no lipid | 63.8 | NA |
| 2 | 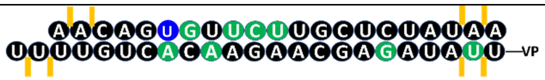 | With 2′-O16 lipid | 62.3 | -1.5 |
| 3 | 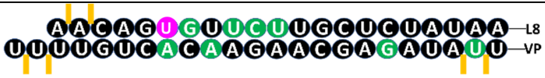 | C5-functionalized-U containing C28 alkyl chain | 55.8 | -8.0 |

[a]2′-fluoro (2′-F) and 2′-*O*-methyl (2′-OMe) nucleotides are indicated in green and black, respectively. Phosphorothioate (PS) linkages are indicated by orange lines. VP refers to vinyl phosphonate. [b]$T_m$ values were obtained from the maxima of the first derivatives of the (average of three heating and three cooling) melting curves ($A$260 vs temperature) recorded in 0.1 × PBS buffer (pH 7.4) using 1.0 $\mu$M concentrations of each strand. [c]$\Delta T_m$ is the difference in melting temperature between the modified duplex and the reference duplex (Entry 1) which had a $T_m$ of 63.8 °C. Cooling curves were recorded from 95 to 25 °C with a cooling rate of 1 °C/min. Heating curves were recorded from 25 to 95 °C with a heating rate of 1 °C/min. The $T_m$ values are accurate within ±1.0 °C.

A

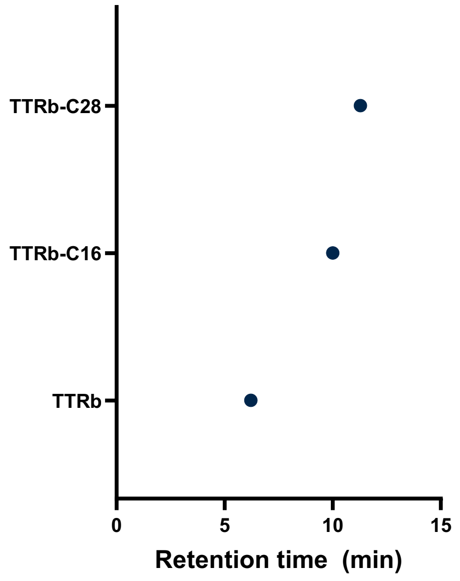


B

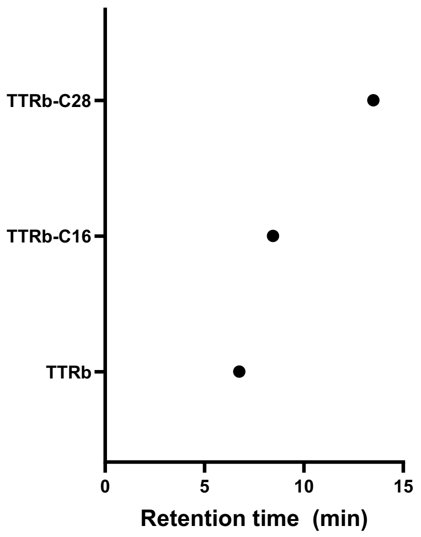


**Figure S2**. Plotted (RP-HPLC) retention time of A) ocTTR duplexes B) ocTTR single strands

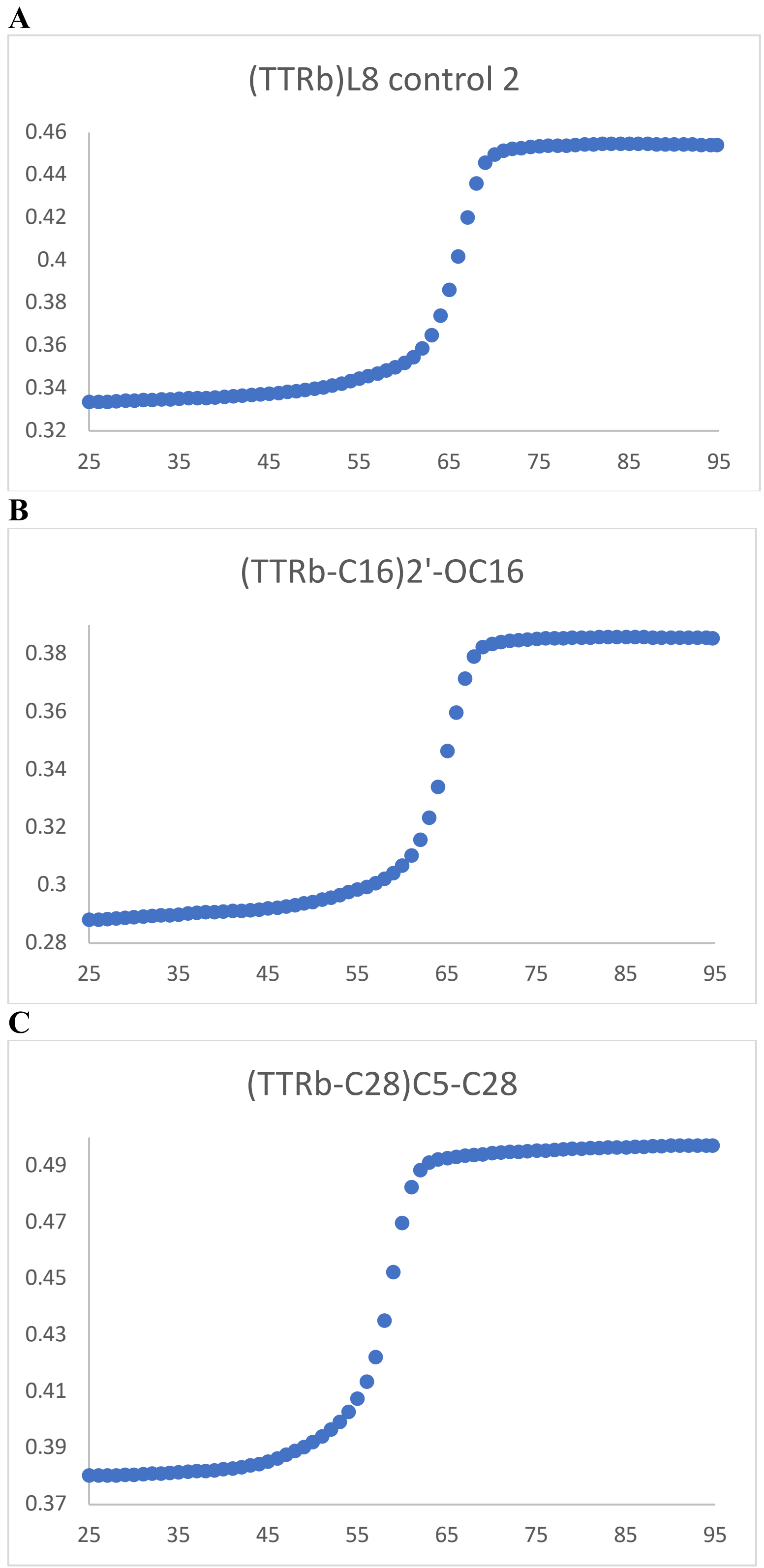


**Figure S3:** Representative heating curves of thermal melting study of A) TTRb B) TTRb-C16 and C) TTRb-C28. Actual $T_m$ in Table S2 is the average of three heating curves and three cooling curves.

**Figure S4**. Analytical data. NMR and HPLC.

$^1$H NMR of **1** (600 MHz) in $CDCl_3$

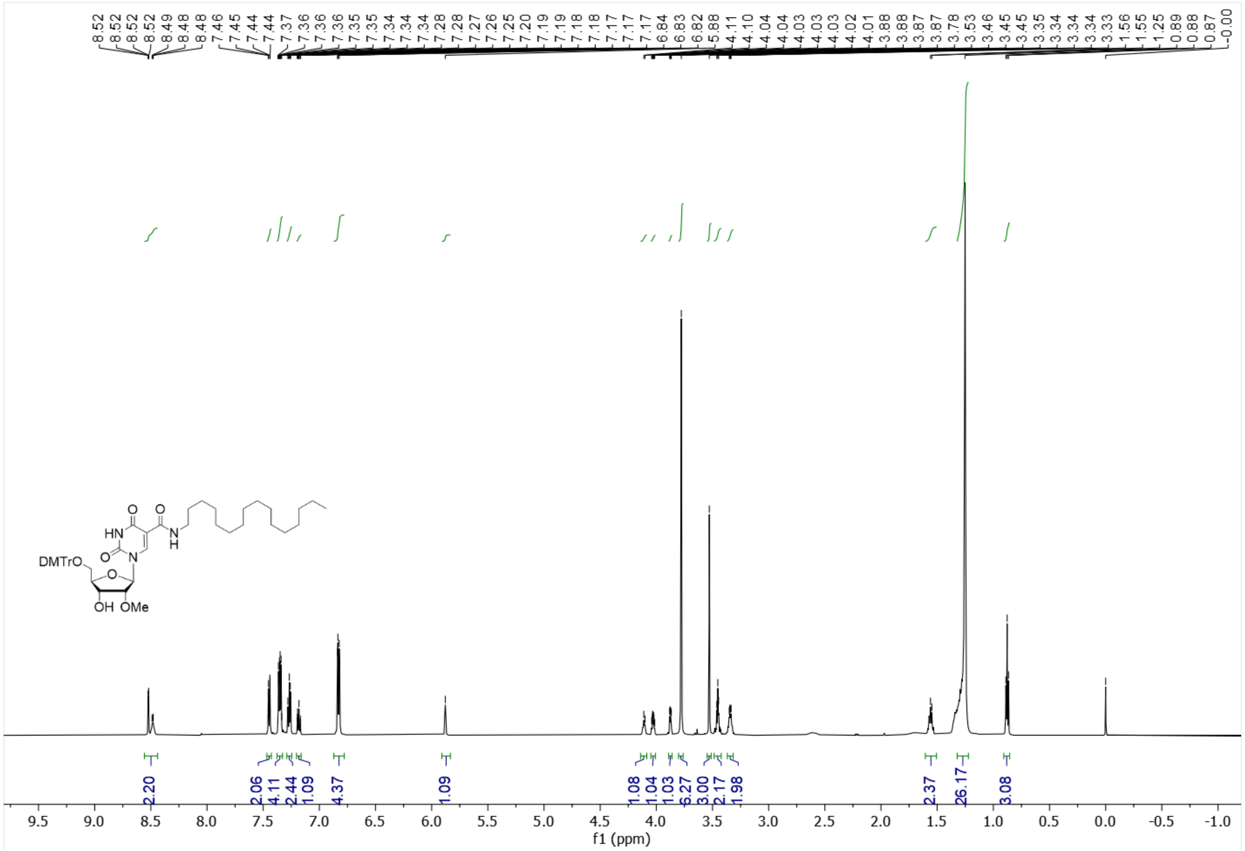

**Figure S4**. Analytical data. NMR and HPLC (cont.)

$^{13}$C NMR of **1** (151 MHz) in $CDCl_3$

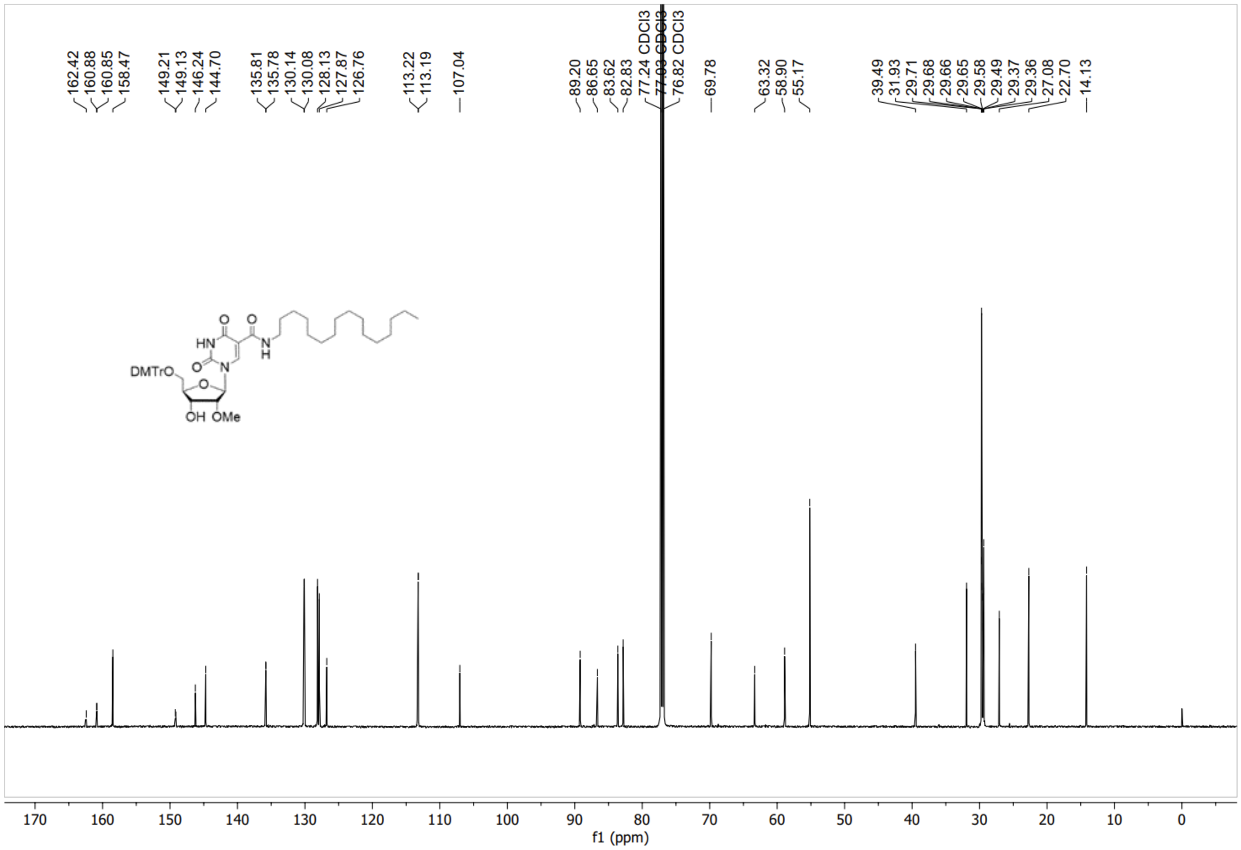

**Figure S4**. Analytical data. NMR and HPLC (cont.)

[1]H NMR of **2** (600 MHz) in $CDCl_3$

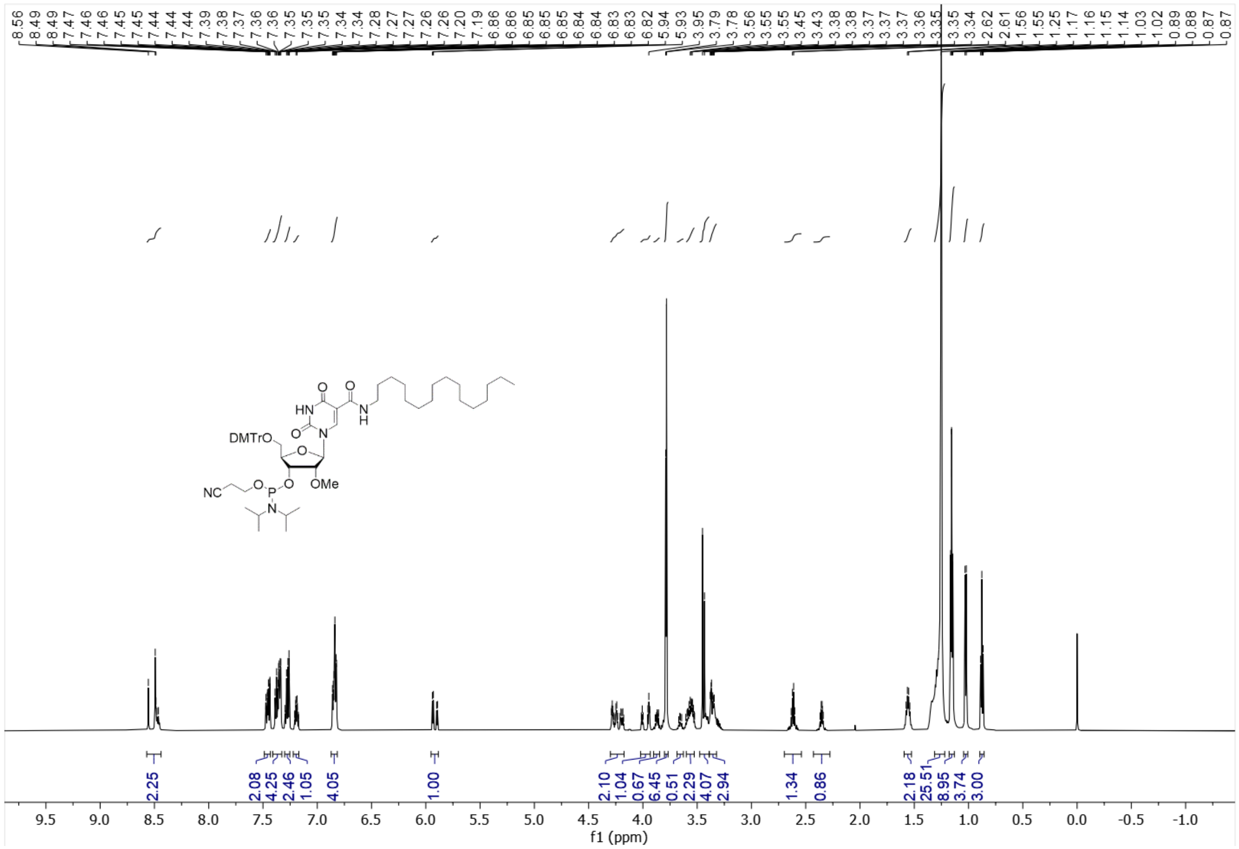

**Figure S4**. Analytical data. NMR and HPLC (cont.)

$^{13}$C NMR of **2** (151 MHz) in $CDCl_3$

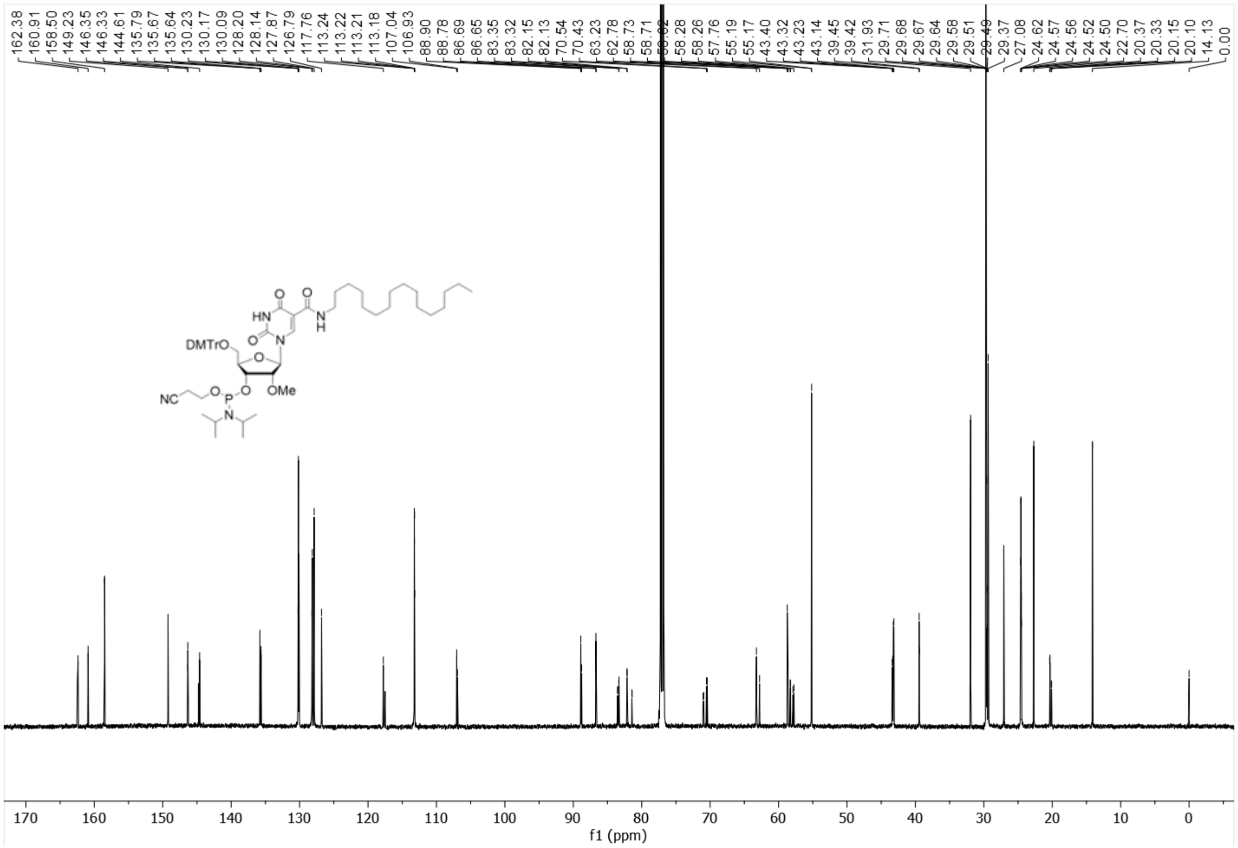

**Figure S4**. Analytical data. NMR and HPLC (cont.)

$^{31}$P NMR of **2** (243 MHz) in $CDCl_3$

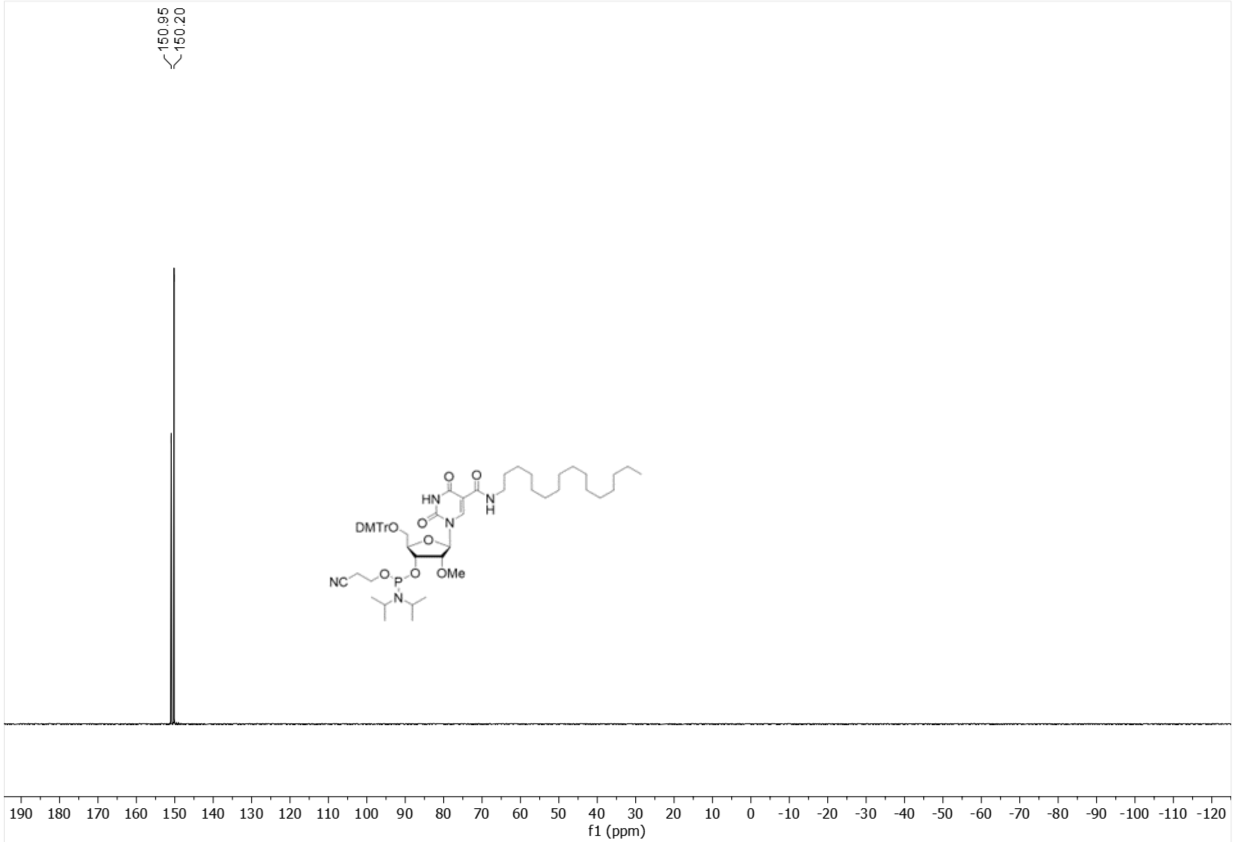

**Figure S4**. Analytical data. NMR and HPLC (cont.)

$^{1}$H NMR of **3** (600 MHz) in $CDCl_3$

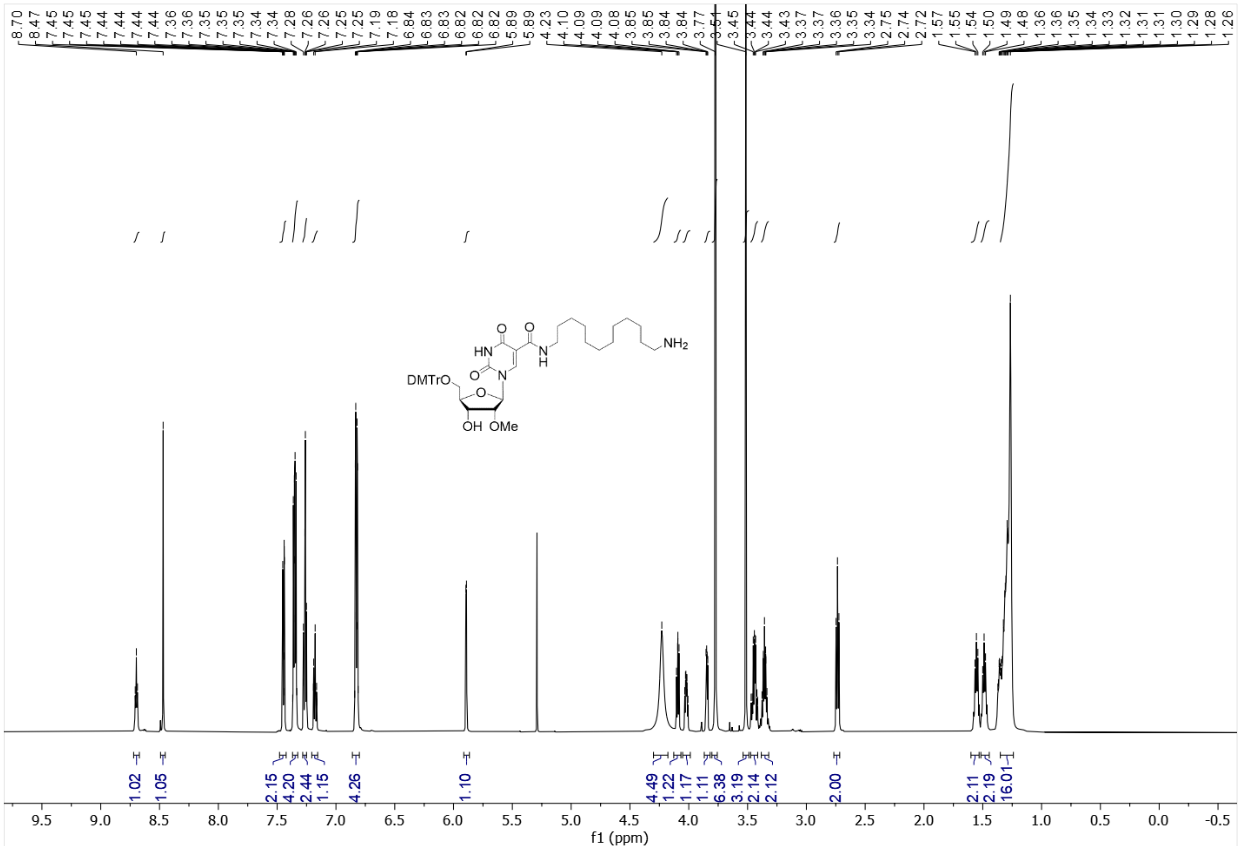

**Figure S4**. Analytical data. NMR and HPLC (cont.)

$^{13}$C NMR of **3** (151 MHz) in $CDCl_3$

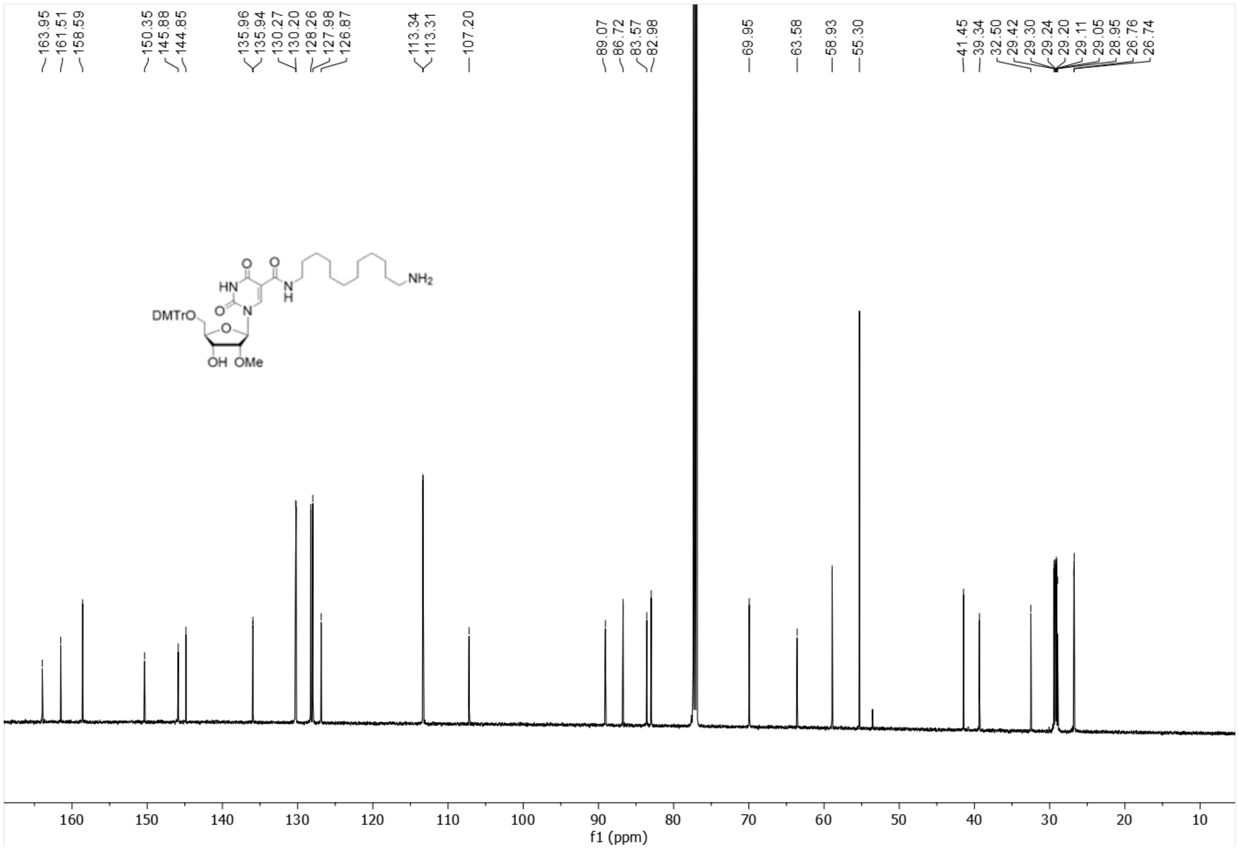

**Figure S4**. Analytical data. NMR and HPLC (cont.)

[1]H NMR of **4**(600 MHz) in $CDCl_3$

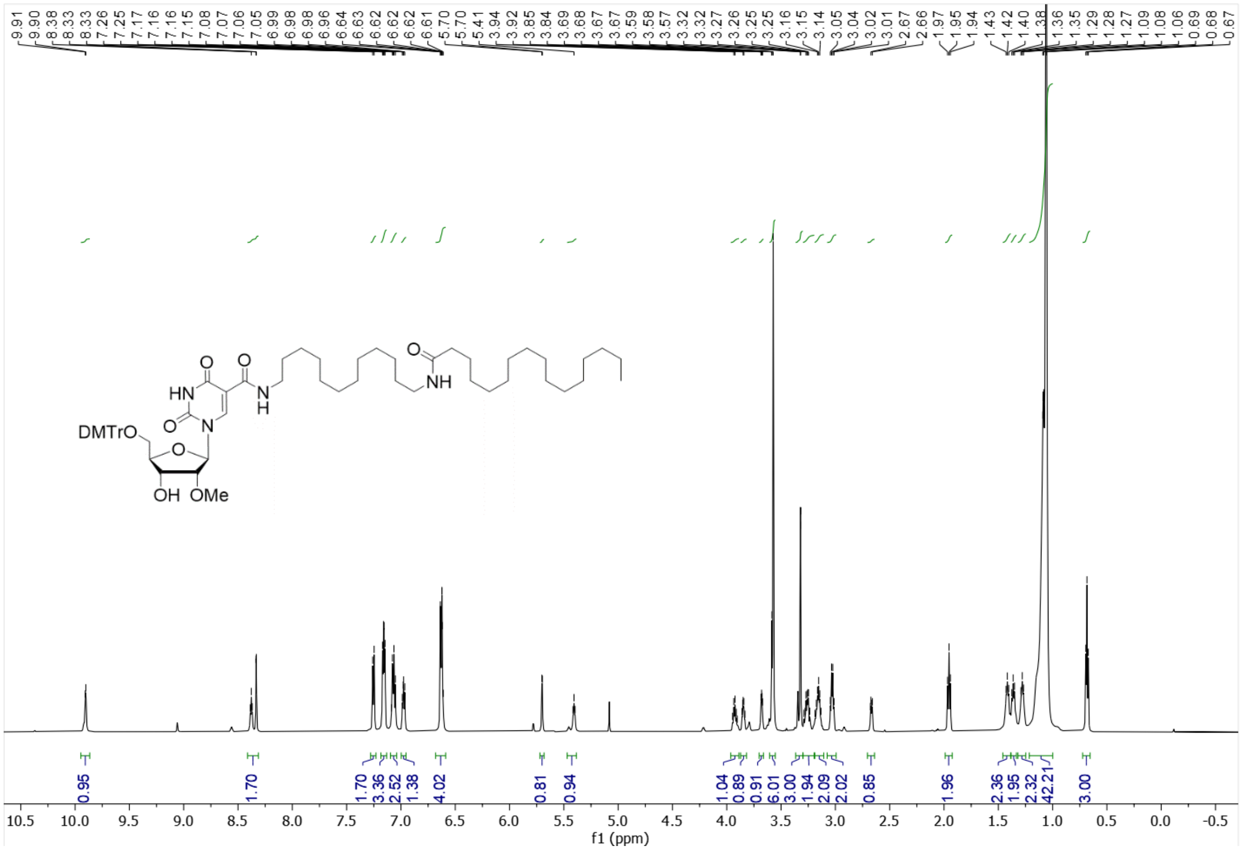

**Figure S4**. Analytical data. NMR and HPLC (cont.)

$^{13}$C NMR of **4** (151 MHz) in $CDCl_3$

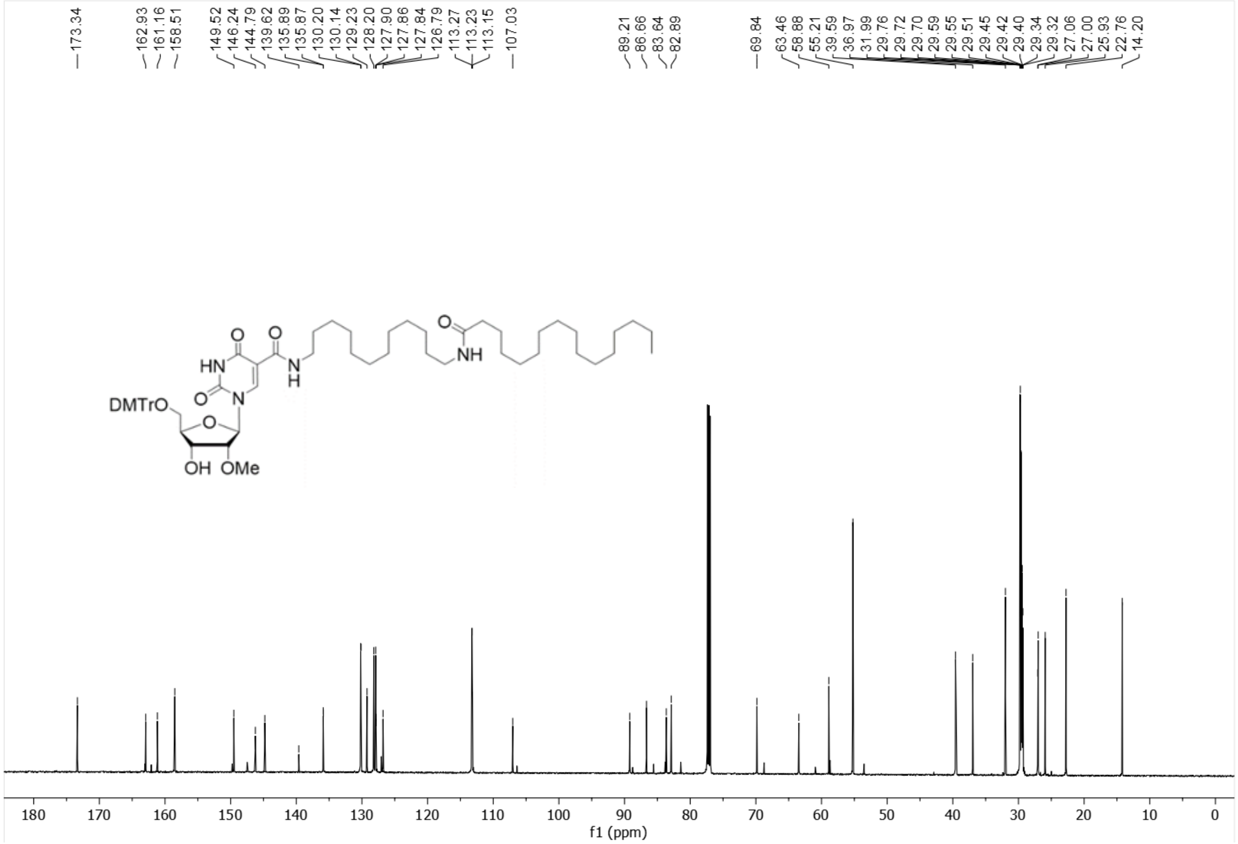

**Figure S4**. Analytical data. NMR and HPLC (cont.)

$^{1}H$ NMR of **5** (600 MHz) in $CDCl_3$

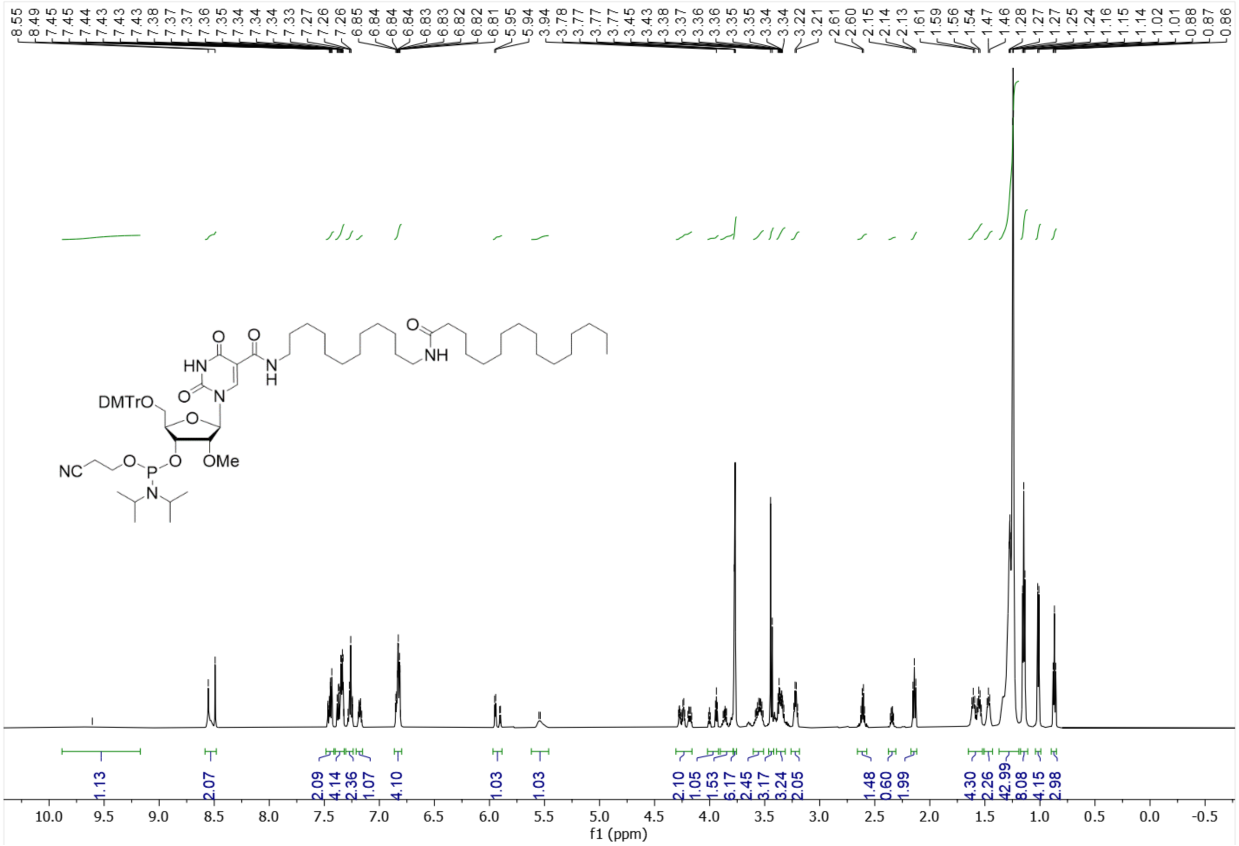

**Figure S4**. Analytical data. NMR and HPLC (cont.)

$^{13}$C NMR of **5** (151 MHz) in $CDCl_3$

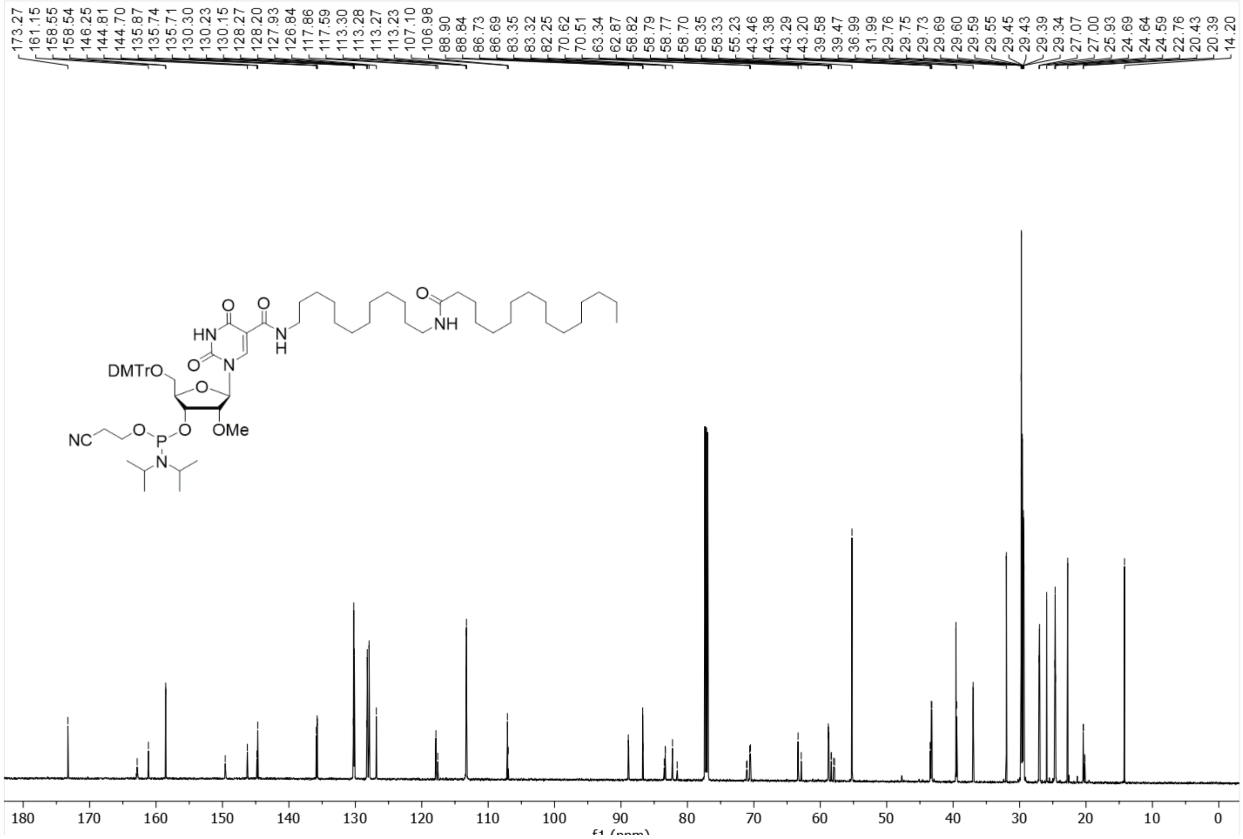

**Figure S4**. Analytical data. NMR and HPLC (cont.)

$^{31}$P NMR of **5** (243 MHz) in $CDCl_3$

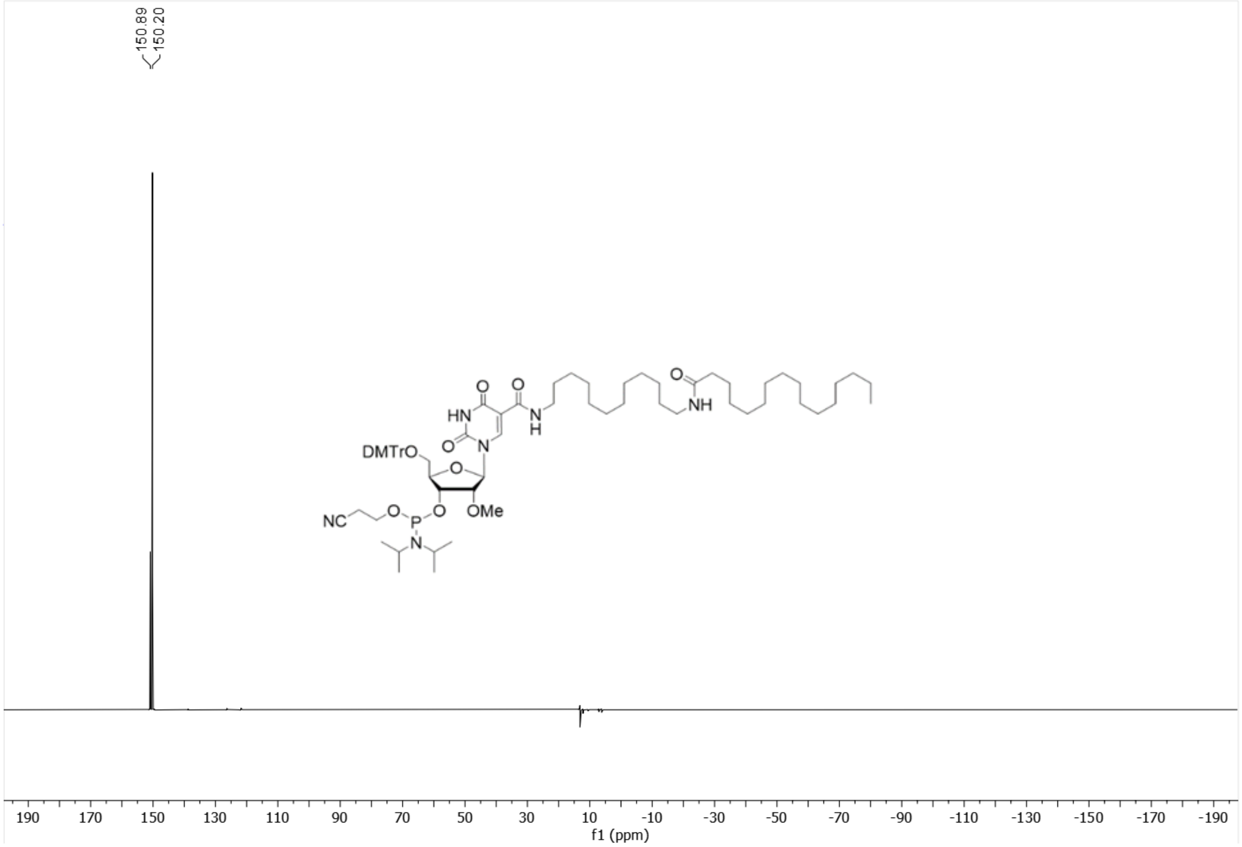

**Figure S4**. Analytical data. NMR and HPLC (cont.)

### HPLC chromatograms of oligonucleotides

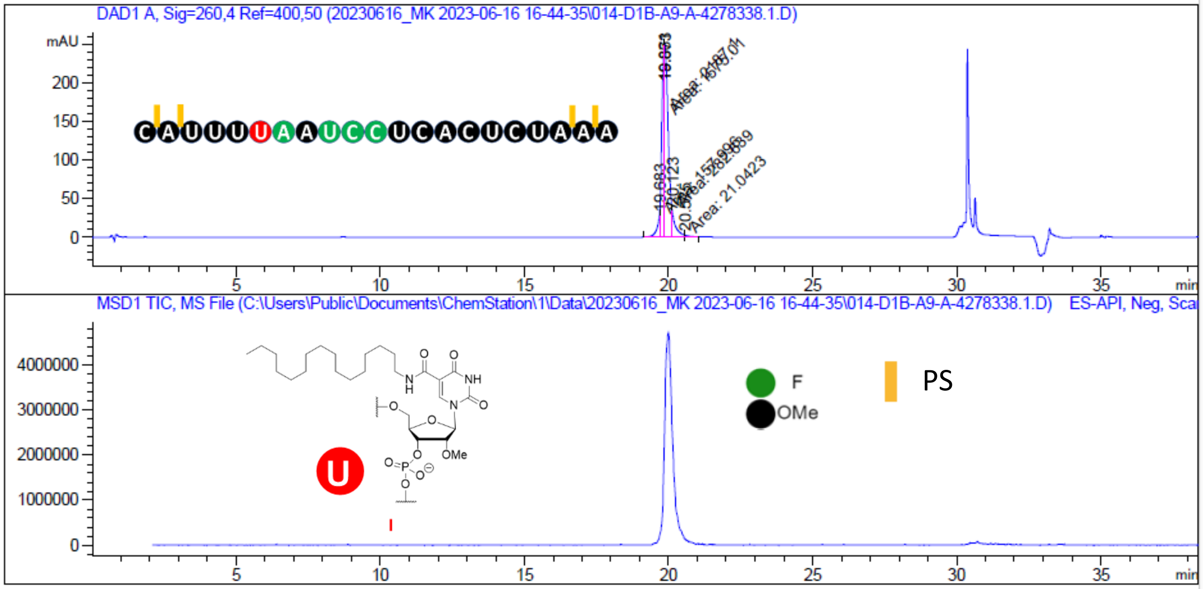


=====================================================================

Area Percent Report

=====================================================================

Sorted By : Signal
Multiplier : 1.0000
Dilution : 1.0000
Do not use Multiplier & Dilution Factor with ISTDs

Signal 1: DAD1 A, Sig=260,4 Ref=400,50

1290 Infinity2 LCMS 1 6/21/2023 3:36:52 PM SYSTEM Page 1 of 2

Data File C:\Users\P...ion\1\Data\20230616_MK 2023-06-16 16-44-35\014-D1B-A9-A-4278338.1.D
Sample Name: A-4278338.1

| Peak # | RetTime [min] | Type | Width [min] | Area [mAU*s] | Height [mAU] | Area % |
|---|---|---|---|---|---|---|
| 1 | 19.683 | MF | 0.0954 | 157.99622 | 27.60995 | 3.7406 |
| 2 | 19.861 | MF | 0.1053 | 1575.00513 | 249.18042 | 37.2890 |
| 3 | 19.873 | MF | 0.1452 | 2187.09937 | 250.99039 | 51.7806 |
| 4 | 20.123 | MF | 0.0986 | 282.63901 | 35.39711 | 6.6916 |
| 5 | 20.575 | FM | 0.1595 | 21.04233 | 2.19864 | 0.4982 |
| Totals : | | | | 4223.78205 | 565.37650 | |

Reverse-phase HPLC profile for **ON1**.

**Figure S4**. Analytical data. NMR and HPLC (cont.)

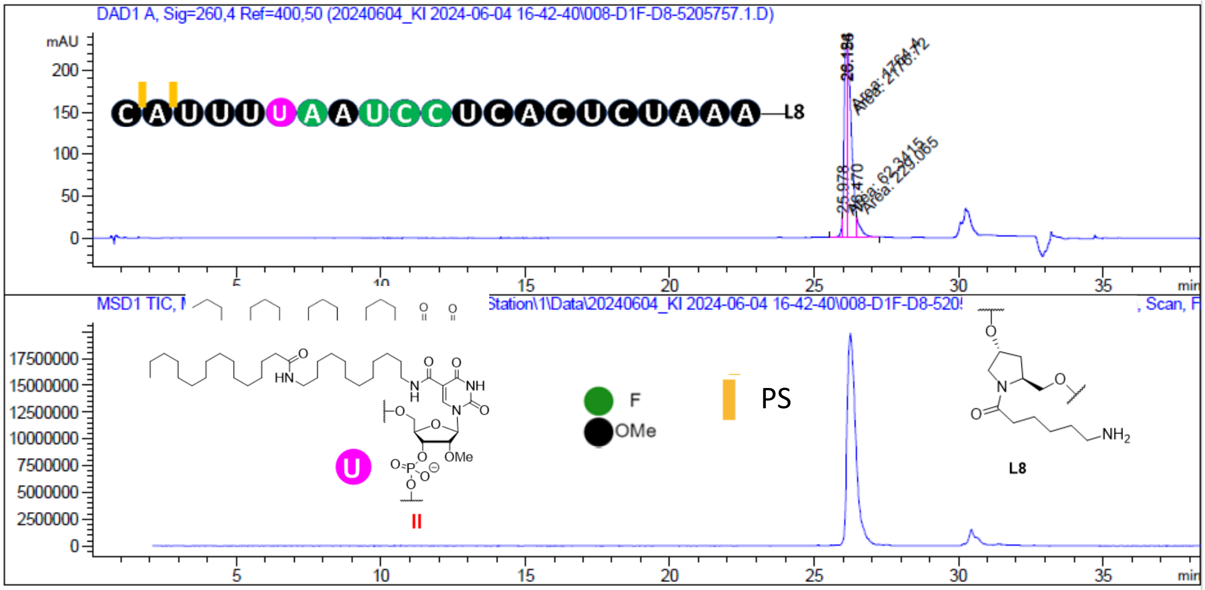


```
=====================================================================
                         Area Percent Report
=====================================================================

Sorted By             :      Signal
Multiplier            :      1.0000
Dilution              :      1.0000
Do not use Multiplier & Dilution Factor with ISTDs


Signal 1: DAD1 A, Sig=260,4 Ref=400,50
```



Data File C:\Users\P...ation\1\Data\20240604_KI 2024-06-04 16-42-40\008-D1F-D8-5205757.1.D
Sample Name: 5205757.1

| Peak # | RetTime [min] | Type | Width [min] | Area [mAU*s] | Height [mAU] | Area % |
|---|---|---|---|---|---|---|
| 1 | 25.978 | MF | 0.0448 | 62.34154 | 23.21166 | 1.4729 |
| 2 | 26.124 | MF | 0.1273 | 1764.40369 | 230.97623 | 41.6868 |
| 3 | 26.156 | FM | 0.1607 | 2176.71704 | 225.81992 | 51.4283 |
| 4 | 26.470 | FM | 0.1534 | 229.06462 | 24.89195 | 5.4120 |
| Totals : | | | | 4232.52689 | 504.89976 | |

**Figure S4**. Analytical data. NMR and HPLC (cont.)

Reverse-phase HPLC profile for **ON2**.

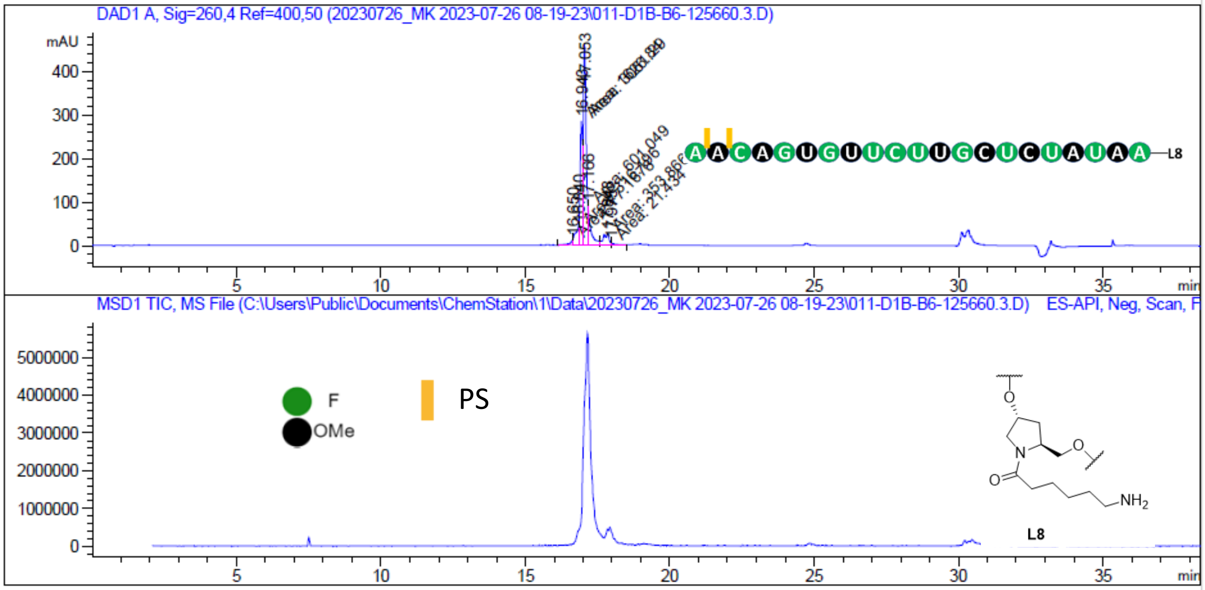


```
====================================================================
                        Area Percent Report
====================================================================

Sorted By              :      Signal
Multiplier             :      1.0000
Dilution               :      1.0000
Do not use Multiplier & Dilution Factor with ISTDs


Signal 1: DAD1 A, Sig=260,4 Ref=400,50
```

1290 Infinity2 LCMS 1 7/26/2023 4:02:26 PM SYSTEM Page 1 of 2

Data File C:\Users\P...tation\1\Data\20230726_MK 2023-07-26 08-19-23\011-D1B-B6-125660.3.D
Sample Name: 125660.3

| Peak # | RetTime [min] | Type | Width [min] | Area [mAU*s] | Height [mAU] | Area % |
|---|---|---|---|---|---|---|
| 1 | 16.650 | MF | 0.1044 | 77.16778 | 12.31973 | 1.2738 |
| 2 | 16.840 | FM | 0.1273 | 316.19632 | 41.40050 | 5.2196 |
| 3 | 16.943 | MF | 0.0954 | 1626.83618 | 284.21198 | 26.8550 |
| 4 | 17.053 | MF | 0.1102 | 3061.29224 | 462.98215 | 50.5344 |
| 5 | 17.166 | MF | 0.0941 | 601.04877 | 89.65131 | 9.9218 |
| 6 | 17.848 | MF | 0.2279 | 353.86646 | 25.87759 | 5.8415 |
| 7 | 17.975 | FM | 0.0890 | 21.43404 | 4.01590 | 0.3538 |
| Totals : | | | | 6057.84178 | 920.45916 | |

**Figure S4**. Analytical data. NMR and HPLC (cont.)

Reverse-phase HPLC profile for **ON3**.

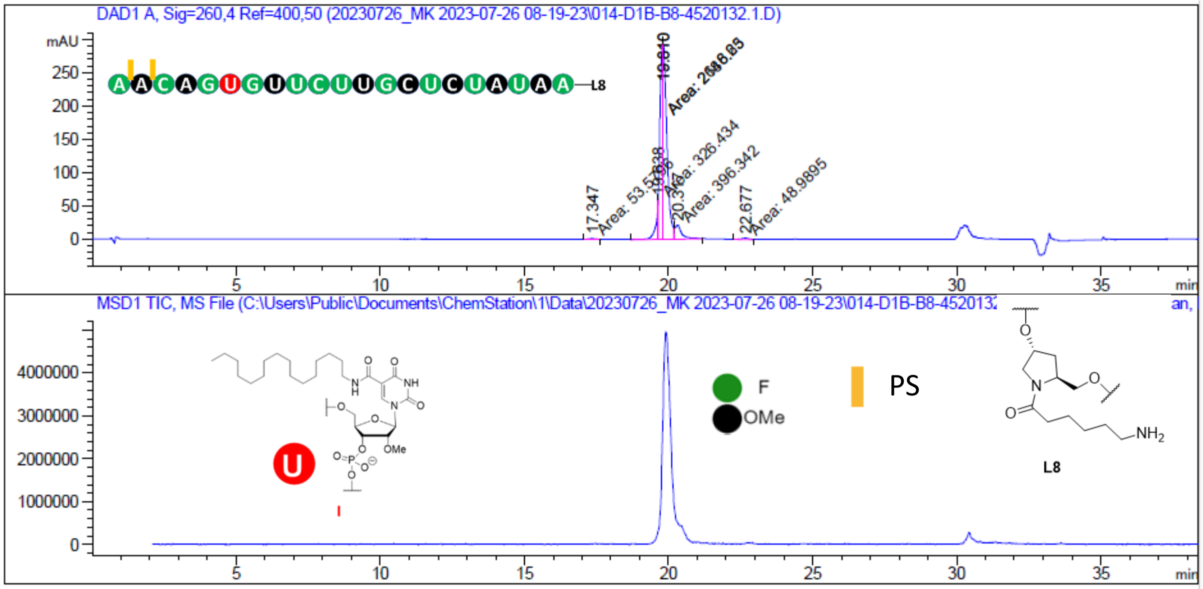


=====================================================================

Area Percent Report

=====================================================================

Sorted By : Signal
Multiplier : 1.0000
Dilution : 1.0000
Do not use Multiplier & Dilution Factor with ISTDs

Signal 1: DAD1 A, Sig=260,4 Ref=400,50

| Peak # | RetTime [min] | Type | Width [min] | Area [mAU*s] | Height [mAU] | Area % |
|---|---|---|---|---|---|---|
| 1 | 17.347 | MM | 0.4097 | 53.57963 | 2.17980 | 0.9466 |
| 2 | 19.638 | MF | 0.0916 | 326.43375 | 59.39338 | 5.7671 |
| 3 | 19.810 | FM | 0.1211 | 2148.62939 | 295.61874 | 37.9601 |
| 4 | 19.810 | FM | 0.1514 | 2686.25342 | 295.63287 | 47.4584 |
| 5 | 20.317 | FM | 0.3142 | 396.34195 | 21.02293 | 7.0022 |

1290 Infinity2 LCMS 1 7/27/2023 12:43:28 PM SYSTEM Page 1 of 2

Data File C:\Users\P...ation\1\Data\20230726_MK 2023-07-26 08-19-23\014-D1B-B8-4520132.1.D
Sample Name: 4520132.1

| Peak # | RetTime [min] | Type | Width [min] | Area [mAU*s] | Height [mAU] | Area % |
|---|---|---|---|---|---|---|
| 6 | 22.677 | MM | 0.3396 | 48.98952 | 2.40406 | 0.8655 |

Totals : 5660.22766 676.25179

**Figure S4**. Analytical data. NMR and HPLC (cont.)

Reverse-phase HPLC profile for **ON4**.

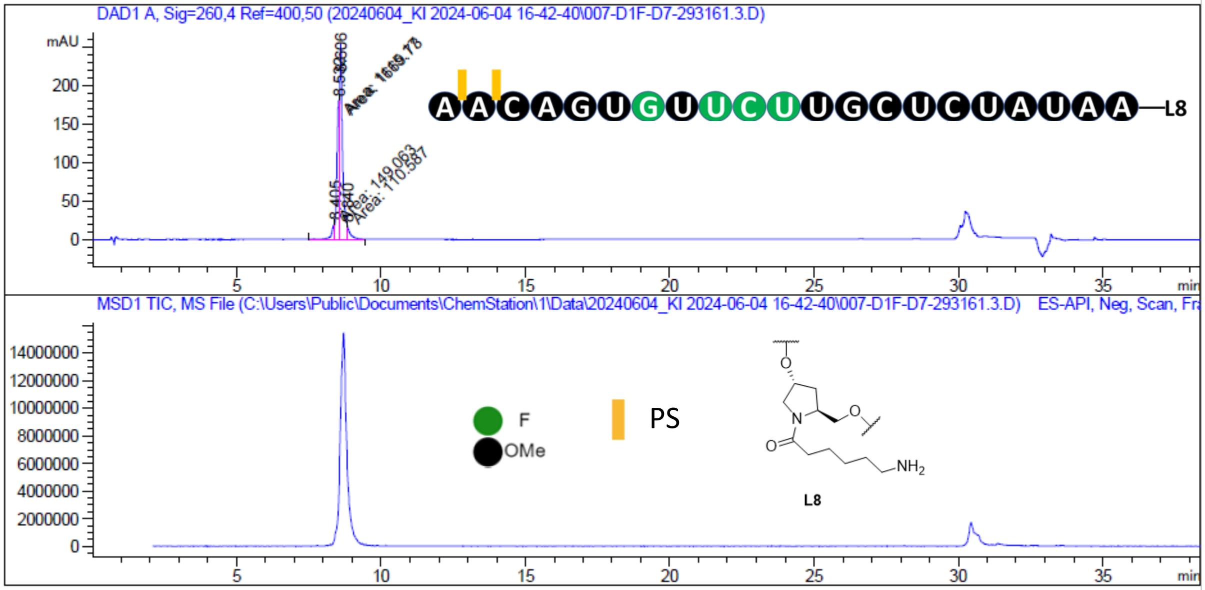


```
=====================================================================
                         Area Percent Report
=====================================================================

Sorted By             :      Signal
Multiplier            :      1.0000
Dilution              :      1.0000
Do not use Multiplier & Dilution Factor with ISTDs


Signal 1: DAD1 A, Sig=260,4 Ref=400,50
```

LCMS_011630 6/5/2024 10:07:57 AM SYSTEM Page 1 of 2

Data File C:\Users\P...tation\1\Data\20240604_KI 2024-06-04 16-42-40\007-D1F-D7-293161.3.D
Sample Name: 293161.3

| Peak # | RetTime [min] | Type | Width [min] | Area [mAU*s] | Height [mAU] | Area % |
|---|---|---|---|---|---|---|
| 1 | 8.405 | MF | 0.1256 | 149.06302 | 19.78739 | 4.8961 |
| 2 | 8.532 | MF | 0.1031 | 1115.17236 | 180.19226 | 36.6285 |
| 3 | 8.606 | FM | 0.1092 | 1669.73010 | 254.80200 | 54.8432 |
| 4 | 8.840 | FM | 0.1108 | 110.58685 | 16.62820 | 3.6323 |

Totals : 3044.55233 471.40985

**Figure S4**. Analytical data. NMR and HPLC (cont.)

Reverse-phase HPLC profile for **ON5**.

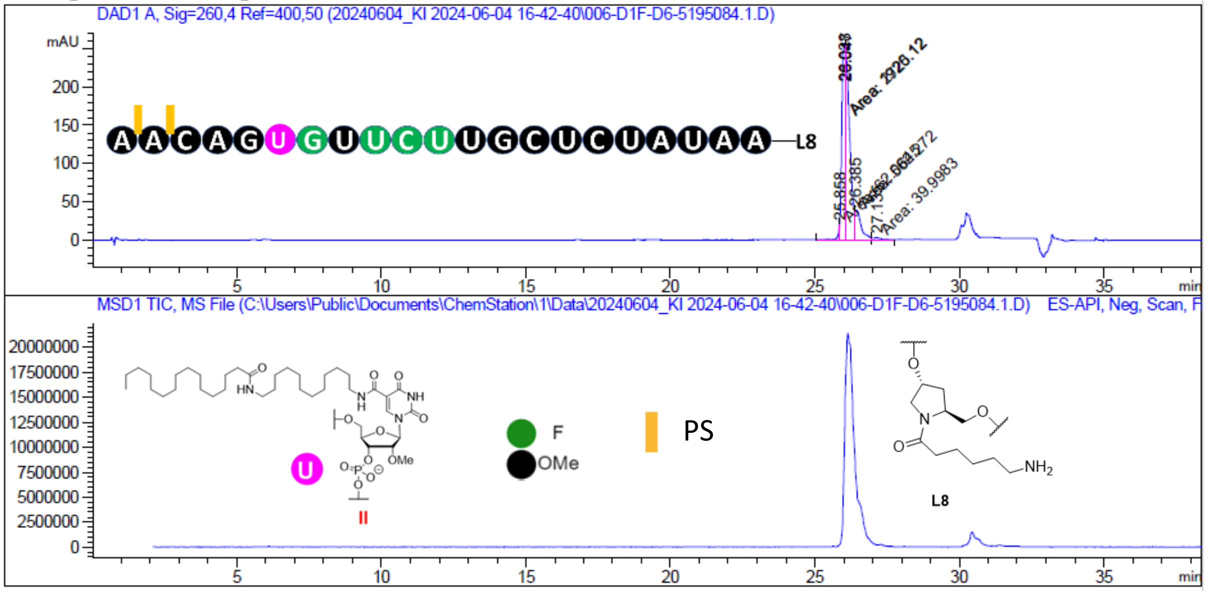


======================================================================

Area Percent Report

======================================================================

Sorted By : Signal

Multiplier : 1.0000

Dilution : 1.0000

Do not use Multiplier & Dilution Factor with ISTDs

Signal 1: DAD1 A, Sig=260,4 Ref=400,50

LCMS_011630 6/5/2024 9:54:49 AM SYSTEM Page 1 of 2

Data File C:\Users\P...ation\1\Data\20240604_KI 2024-06-04 16-42-40\006-D1F-D6-5195084.1.D

Sample Name: 5195084.1

| Peak # | RetTime [min] | Type | Width [min] | Area [mAU*s] | Height [mAU] | Area % |
|---|---|---|---|---|---|---|
| 1 | 25.858 | MF | 0.0517 | 62.06452 | 20.01926 | 1.1689 |
| 2 | 26.028 | MF | 0.1251 | 1920.12292 | 255.85619 | 36.1634 |
| 3 | 26.047 | FM | 0.1789 | 2725.11841 | 253.92242 | 51.3246 |
| 4 | 26.385 | MF | 0.1779 | 562.27197 | 38.16459 | 10.5898 |
| 5 | 27.135 | FM | 0.2719 | 39.99833 | 2.45222 | 0.7533 |

Totals : 5309.57616 570.41468

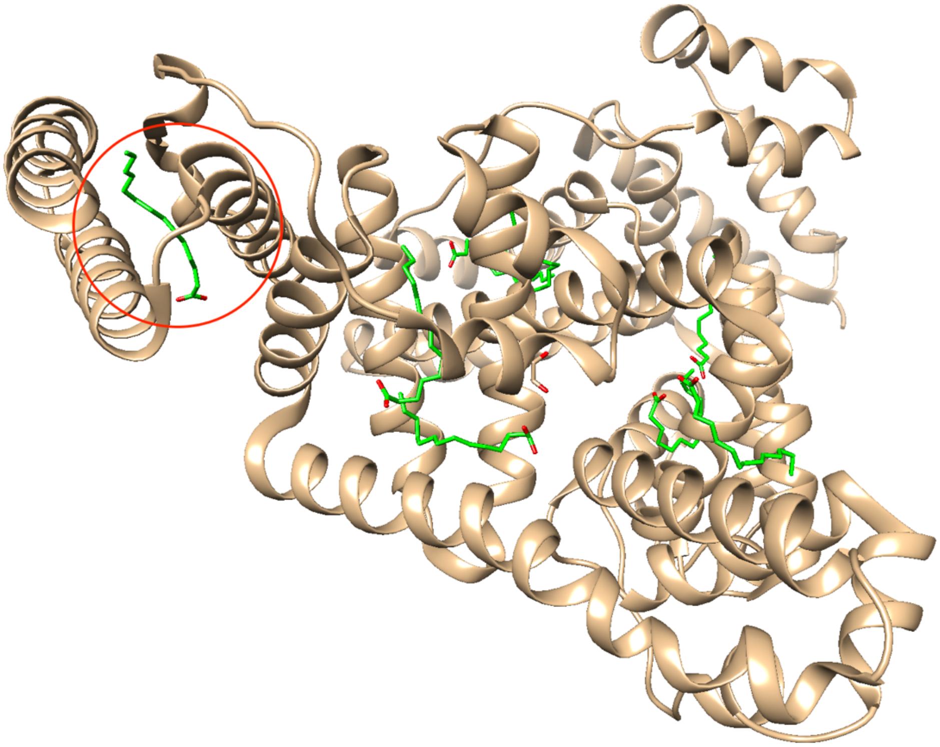

**Figure S5.** Crystal structure of human serum albumin in complex with myristic acid (PDB ID 8RCP). The protein is shown in cartoon mode and colored in tan and carbon atoms of eight bound fatty acid molecules are colored in green. We picked the myristic acid molecule marked with a red circle for building the TTRb-O2'-C16 and TTRb-C5-C28 siRNA duplexes bound to albumin.
